# Overcoming Immunological Resistance Enhances the Efficacy of a Novel anti-tMUC1 CAR T Cell Treatment Against Pancreatic Ductal Adenocarcinoma

**DOI:** 10.1101/642934

**Authors:** Mahboubeh Yazdanifar, Ru Zhou, Priyanka Grover, Chandra Williams, Mukulika Bose, Laura Jeffords Moore, Shu-ta Wu, Richard Chi, John Maher, Didier Dreau, Pinku Mukherjee

**Author notes:** corresponding author, Pinku Mukherjee.

## Abstract

Chimeric antigen receptor engineered T cells (CAR T cells) have shown remarkable success in treating hematologic cancers. However, this efficacy has yet to translate to treatment in solid tumors. Pancreatic ductal adenocarcinoma (PDA) is a fatal malignancy with poor prognosis. Treatment options are limited and commonly associated with severe side effects. We have developed and characterized a second generation CAR engineered T cell using the light and heavy chain sequence derived from a novel monoclonal antibody, TAB004, that specifically binds the tumor associated antigen, tMUC1. tMUC1 is overexpressed in varying levels on ~85% of all human PDA. We present data showing that the TAB004 derived CAR T cells (tMUC1-CAR T cells) specifically bind to tMUC1 on PDA cells and is cytotoxic against the majority of the PDA cell lines. The tMUC1-CAR T cells do not bind or kill normal epithelial cells. We further demonstrate that the tMUC1-CAR T cells control the growth of orthotopic pancreatic tumors *in vivo.* PDAs are generally cold tumors with resistance to many standard treatment modalities, thus, it was not surprising that some of the PDA cell lines were refractory to CAR T cell treatment. qPCR analysis of several genes known to be associated with immune resistance revealed overexpression of indoleamine 2, 3-dioxygenases-1 (IDO1), Cyclooxygenase 1 and 2 (COX1 and COX2), Adenosine deaminases acting on RNA (ADAR1) and galectin-9 (Gal-9). We treated resistant PDA cells with a combination of CAR T cells and biological inhibitors of IDO1, COX1/2, ADAR1, and Gal-9. Results showed a significant enhancement of CAR T cell cytotoxicity against PDA cells when inhibiting IDO1, COX1/2, and Gal-9 but not ADAR1 or COX2. Overcoming CAR T cell resistance in PDA is a significant advancement in the field and may lead to future combination therapies that are less toxic but more efficient against this deadly disease.

## Introduction

Pancreatic Ductal Adenocarcinoma (PDA) which arises from exocrine cells of the pancreas [1] is one of the deadliest forms of cancer with mortality rate closely parallel to incidence [2]. It is the third leading cause of cancer related deaths in the United States, yet treatment options are limited and often associated with a high recurrence rate and poor prognosis [3]. PDA, once diagnosed is already resistant or soon become resistant to conventional therapies [4]. Thus, developing new strategies targeting resistant PDA is warranted.

Chimeric antigen receptor T cell (CAR T cell) therapy is an exciting approach that arm T cells with a chimeric receptor that can recognize a surface antigen on tumor cells [5]. CAR T cell therapy has shown enormous success in treating hematologic malignancies [6, 7] and metastatic melanoma [8]; however, this success has not been extended to adenocarcinomas [9]. This may be due to the limited selection of antigens that are expressed at a high level on the surface of solid tumors.

MUC1, a transmembrane glycoprotein expressed at the apical surface of epithelial cells [10] [11] is recognized as the second most targetable antigen by the National Cancer Institute [12]. The aberrant tumor-associated form of MUC1 (tMUC1) is predominantly expressed in >80% of human PDA [13] [14] and is a key modulator of several signaling pathways that affect oncogenesis, motility, and metastasis. tMUC1 overexpression occurs at the early stages of the disease [15] and high expression is associated with poor prognosis in PDA patients [16].

Few studies have tested CAR T cell therapy in PDA targeting antigens such as mesothelin [17], NY-ESO-1 (NCT01967823) and ROR1 [18] [19]. MUC1 has also been targeted in CAR T cell settings by our collaborators [20–22] and two other groups [23] [24] and shown promising results, but needs further validation. Most of CAR T cell clinical trial on PDA have targeted mesothelin, which is also expressed on the normal mesothelial cells. There is only one clinical trial investigating an anti-MUC1 CAR T cell for treating patients with MUC1 positive advanced refractory solid tumors including PDA (NCT02587689) [25]. The result of this trial shows adverse reactions in patients. This may be due to using scFv from an anti-MUC1 antibody (Ab) that also reacts with normal MUC1 expressed on normal cells.

In this study, we introduce a novel anti-tMUC1 CAR T cell using scFv derived from a highly specific anti-tMUC1 monoclonal Ab, TAB004, that does not recognize the normal form of MUC1 [26, 27]. TAB004 detects tMUC1 on tissues of PDA and breast cancer patients, on PDA cells, and PDA stem cells but spares recognition of normal epithelial cells [28–30]. Previously, we have shown specific localization of TAB004 in breast and pancreatic tumors in human MUC1 transgenic mouse models (in which the entire glandular epithelia express normal human MUC1) further validating the tumor-specificity of TAB004 [27, 31, 32]. This specificity is critical, since targeting shared antigen by CAR T cells may cause fatal toxicity [33] [34]. We tested the engineered tMUC1-CAR T cells against a panel of human PDA cell lines with varying expression levels of tMUC1 as well as against normal epithelial cells and fibroblasts.

PDA is an immunologically cold tumor with known resistance to a variety of therapies including immunotherapy [35]. Thus, it was not surprising that we found that some of PDA cells were refractory to tMUC1-CAR T cell treatment. Gene expression profile of resistant vs. sensitive PDA cells revealed the overexpression of several genes associated with immune tolerance such as IDO1, COX1/2, ADAR1, and Gal-9. Thus, we hypothesized that blocking the above proteins can overcome the immune tolerance of resistant PDA cells to tMUC1-CAR T cell therapy. We present data indicating that combination therapy of tMUC1-CAR T cells with biological inhibitors of resistance inducing genes resulted in significant enhancement of tMUC1-CAR T cell cytotoxicity against resistant PDA cells. These novel combinations may have major clinical significance in designing future CAR T cell therapies against resistant PDA.

## Materials and methods

### Cells

All the PDA cell lines used in this study were originally purchased from American Type Culture Collection (ATCC, Manassas, VA 20110, USA) and cultured as instructed. Primary T cells were obtained from human peripheral blood mononuclear cells (PBMCs), which were bought from STEMCELL Technologies (Cambridge, MA #70025.1). Normal human fibroblasts and breast epithelial cells were obtained from Coriell Institute (NJ, USA). The BxPC3-MUC1 cell line was made by retroviral transfection of BxPC3 wild-type (ATCC) with the PLNCX.1 plasmid (Mayo Clinic K1060-C), which contains the full length human MUC1 gene. BxPC3-Neo was transfected with the empty vector PLNCX.1 plasmid. The MiaPaCa2-Luc cell line was generated by Lipofectamine transfection (Lipofectamine 3000, Invitrogen, #L3000015) of the MiaPaCa2 cell line with pGL4.50[luc2/CMV/Hygro] vector (Promega).

### CAR constructs and cloning

A second generation anti-MUC1 CAR harboring TAB004 Ab scFv was synthesized by subcloning the scFv from TAB004 [26] into the SFG-based retroviral backbone plasmid encoding the transmembrane and intracellular domains of CD28 and CD3ζ (synthesized by Dr. John Maher’s group [36]). This CAR contains a myc tag for detection. The CAR-mKate construct was made by cloning PCR-amplified tMUC1-CAR sequence into the PLNCX.1 retroviral vector (Mayo Clinic K1060-C) at the MluI site along with the mKate2 sequence fused through a GA linker. The mKate2 gene (pFA6a-mkate-kanmx6) was a gift from Dr. Richard Chi. CTL-CAR was created by PCR cloning of CAR in three fragments missing the majority of the TAB scFv sequence. All cloning was done using NEBuilder^®^ HiFi DNA Assembly Cloning Kit (NEB #E5520). PCRs were done using Q5^®^ High-Fidelity DNA Polymerase (NEB #M0491).

### Viral transfection of T cells

Retroviruses were made by transduction of GP2-293 packaging cell line with 10ug CAR DNA and pVSV envelope plasmid. 48 hrs viral supernatant was used for infection of T cells. Human PBMCs were activated by CD3/CD28 beads (Dynabeads, Gibco #111.61D) 3 days prior to infection. Non-tissue culture plates (Corning #351146) were coated with retronectin (1mg/ml) (Takara, Mountain View, CA) and incubated at 4 °C at least 12 hrs before infection. The following day, retronectin was removed and the plates were blocked with 2% BSA for 30 minutes. Then, viral supernatant was applied to retronectin-coated plates, which were subsequently centrifuged at 2000g for 2 hrs at 32 °C. Viral supernatants were removed and activated T cells were added to the coated plates. Plates with cells were spun at 1000g, at 32 °C for 10 min and incubated overnight. Cultures were maintained in complete RPMI with 200-300 U/ml human recombinant IL-2 (PeproTech, Rocky Hill, NJ #200-02) and media was refreshed every 3 days. To avoid T cell exhaustion, from day 10 onward cultures were maintained at 50 U/ml IL-7 and IL-15 (PeproTech, Rocky Hill, NJ #200-07). T cells that had been activated but not transduced were used as mock T control. CAR expression level was characterized by flowcytometry using anti-myc tag Ab, and T cells were used between days 11 to 20 after infection.

### T cell cytotoxicity

To test T cell cytotoxicity against target cells, 5,000-10,000 cancer cells or normal cells were plated in triplicate in 96 well plates one day prior to co-culture. Mock or CAR T cells were counted and added to cancer cells at the indicated target to effector (T:E) ratio. Cell viability was evaluated by MTT assay (MTT 500 ug/ml, Sigma) 24, 48, and 72 hrs after co-culture according to the product instructions. The OD value at 540 nm was read and percentage survival was calculated as 100-[(mock T OD – CAR T OD)/mock T OD × 100]. To measure the spontaneous cytotoxicity of T cells, an LDH-based technique, CytoTox 96^®^ Non-Radioactive Cytotoxicity Assay (Promega, USA) was used. BxPC3-Neo and BxPC3-MUC1 cells were plated in 96 well plate (20,000 cells/well) and incubated with 200,000 mock or CAR T cells (T:E 1:10) for 8, 16 and 24 hrs. The amount of released LDH and subsequent cytotoxicity was measured and calculated according to the manufacturer’s instructions.

### Binding assay

HPAFII, a moderate-to-high MUC1 cancer cell line, was plated in 6 well plates (150,000 cells/well) and incubated at 37 °C overnight. Next, the cells were stained with nuclei live cell stain Hoechst (Thermo Fisher #33342) for 30 minutes and washed 3X. 1×10^6^ CAR T cells or CTL T cells (both expressing CAR constructs fused to mKate fluorescent tag) were added to the respective wells and plates were incubated at 37 °C for 4 hrs with occasional rocking. Afterward, media was removed and cells were washed 2X and then imaged by DeltaVision workstation (Applied Precision).

### Flowcytometry

The CAR expression level was quantified using myc tag-FITC staining (Cell Signaling Technology, Danvers, MA). T cell subtypes were determined by staining for CD4-PE/Cy7 and CD8-eF450 (BD Biosciences, San Jose, CA). PD1-APC (eBioJ105), PDL1-FITC (clone MIH2), IFN-γ-APC (clone 4S.B3) and perforin-PE (clone dG9) Abs were obtained from eBioscience. The human tMUC1 expression on PDA cells was measured by staining with TAB004 primary Ab (provided by OncoTab Inc., Charlotte, NC) and FITC-anti-mouse secondary Ab (Invitrogen #31535). Dead cells were excluded by 7-AAD staining (BD Biosciences #555816). Data were acquired on BD LSRFortessa flowcytometer (BD Biosciences), and analyzed with FlowJo software (version 8.8.7, Tree Star Inc).

### ELISA

The supernatant of PDA cells co-cultured with T cells was assessed for released IFN-γ and granzyme B after 72 hrs of co-culture, using Human IFN gamma Uncoated ELISA Kit (Life Technologies 88-7316-22) and Human Granzyme B DuoSet ELISA (R&D DY2906-05). Shed MUC1 was measured using human MUC1 ELISA kit (OncoTAb Inc., Charlotte, NC). All of the ELISAs were performed on the supernatant after 72 hrs of co-culture.

### RT-PCR, qPCR

RNA was extracted from cancer cells by RNeasy Plus Mini Kit (Qiagen #74134). RT-PCR was performed using AccessQuick™ RT-PCR system (Promega #A1700) and samples were ran on 1.2% agarose gel. The human MUC1 primers used were: Forward TGC ATC AGG CTC AGC TTC A, Reverse GAA ATG GCA CAT CAC TCA G, and Tm 60 °C. qPCR primers were designed using the NCBI primer design tool and synthesized by MWG Eurofins (Louisville, USA). The primer sequences will be available upon request. The relative expression levels of multiple genes such as MUC1, PD1, PDL1, LIF, VEGF, IDO1/2, COX1/2, ADAR1, TGFB, TGFBRI/II, PDGF, M6PR, Gal-9 and TRAIL were quantified in cancer cells before and after exposure to mock and CAR T cells using Applied Biosystems^®^ 7500 fast Real-Time PCR machine and SYBR Green PowerUp Master Mix (Life Tech, A25742).

### Apoptosis assay

Mock and CAR T cells were stained with Annexin V/propidium iodide (PI) dyes according to the Dead Cell Apoptosis Kit protocol (Life Technology V13242) before and after co-culture with cancer cells for 24, 48, and 72 hrs. The percentage of positive cells was assessed using flowcytometry.

### Proliferation assay

HPAFII, CFPAC and MiaPaCa2 cells were plated in 6w plate (500,000 cells/well). Next day, mock and CAR T cells were added to the cancer cells at 1:5 T:E ratio. Number of live T cells per well were calculated at 24, 48 and 72 hrs using trypan blue (0.4%, Thermo Fisher) staining and Countess II automated cell counter (Life Technologies, Carlsbad, USA).

### Imaging

1×10^6^ CAR T cells expressing CAR-mKate were plated in 35 mm Poly-d-lysine Coated MatTek dish (MatTek #P35GCOL-0-14-C) for 24 hrs. The next day, cells were stained with nuclei live cell stain, Hoechst (Thermo Fisher #33342) for 30 minutes and washed gently once, then imaged by DeltaVision workstation (Applied Precision). Image analysis was done using Softworx 6.1 (Applied Precision Instruments).

Videos of CAR T cells killing BxPC3-MUC1 vs. BxPC3-Neo cells were taken by time lapse imaging using a DeltaVision OMX-SR imaging system (GE #29115476). Cancer cells and CAR T cells were co-cultured in 35 mm MatTek dish and placed in the microscope’s 37 °C 5% CO_2_ incubator overnight. Images were taken over the course of 8 hrs at 7 min intervals. PI dye was added to the culture at 50 ng/ml [37]. Only apoptotic cells are susceptible to PI infusion, and turn red upon absorption. Pictures were analyzed using ImageJ v1.51f program (Rasband).

### Combination therapy with drugs and blocking antibody

To assess the effect of drugs on CAR T cell cytotoxicity, cancer cells were plated at 5,000/well concentration in 96 well plates in triplicate. After 24 hrs, the cells were treated with drugs (1-MT [Sigma, #452483], indomethacin [Sigma, #17378], gemcitabine hydrochloride [Sigma, #G6423], paclitaxel [Invitrogen, Taxol equivalent, #P3456] and 5-Fluorouracil [Sigma, F6627]) at indicated concentrations for 24 hrs. Then, drugs were removed and cells were incubated with mock or CAR T cells at T:E ratio of 1:10 for 72 hrs. Next, the percentage survival of cancer cells was measured using MTT assay. For Gal-9 and PD-1 blocking experiments, cancer cells were plated at 10,000 cells/well concentration in 96 well plates in triplicate. After 24 hrs, mock or CAR T cells were added at T:E ratio of 1:10. 0.1, 1, and 10 ug/ml of Gal-9 (clone 9M1-3, BioLegend) or PD-1 blocking Ab (clone EH12.2H7, BioLegend) were added to the media and incubated at 37 °C for 72 hrs. Survival level was measured using MTT assay.

### Animal study

12 female Non-obese diabetic (NOD)-SCID gamma (NOD.Cg-prkdcscidIl2rgtm1w1; NSG™, Jackson Laboratory) mice were anesthetized. Using aseptic techniques, an incision was made in abdominal area off midline just above the pancreas. Then pancreas was gently retracted and injected with 0.5×10^6^ MiaPaCa2-Luc cancer cells. The abdominal incision was sutured and the skin layers were closed using surgical clips. Mice were monitored and their body weight were checked daily for a week. 7 days post-surgery, tumor presence was confirmed using *in vivo* imaging system (IVIS, PerkinElmer). On day 8 post-surgery, when average ROI for pancreatic tumors was 1.4×10^6^, mice were randomized into two groups (with even distribution of tumor size) based on their baseline luminescence intensity. One group received intravenous (IV) injection of Mock T (n=6) and the other received tMUC1-CAR T cells (n=6) (10×10^6^ per mouse). Mice were imaged weekly by IVIS using chemo-luminescence, open filter setting in Living Image 4.3 software. On day 68 post injection, mice were sacrificed and tumors were harvested and weighted. Two mice (1 per group) died of irrelevant cause before the endpoints. To assess T cell trafficking in mice after injection, mock or CAR T cells were labeled with Vivotrack-680 dye (PerkinElmer) according to the manufacturer’s protocol. Six MiaPaCa2-Luc tumor-bearing NSG mice were injected with either 4×10^6^ labeled CAR T or mock T cells through tail vein (n=3). Mice were sacrificed 24 hrs after injection and tumors were dissected and imaged by IVIS. Images were acquired using fluorescence Vivotrack-680 channel with excitation=676 and emission=696 nm, and analyzed using Living Image 4.3 software. Our animal studies were approved by the Institutional Animal Care and Use Committee of the University of North Carolina at Charlotte. All the experimental procedures complied with institutional guidelines.

### Statistical analysis

All of the data were analyzed by Prism (version 8.0; GraphPad Software) and results were presented as mean ±S EM. Data are representative of two or more independent experiments. The statistical analysis was done using Prism software and significance was determined using unpaired Student’s t-test, two-way ANOVA, or Non-parametric Mann-Whitney U test where indicated (*, p< 0.05, **, p< 0.01, ***, p< 0.001).

## Results

### CAR architecture, CAR expression on engineered T cells, and binding of CAR T cells to target PDA cells

The architecture of CAR constructs used in this study is illustrated in figure 1A. TAB004 Ab’s variable fragments are cloned into a 2^nd^ generation CAR plasmid (SFG muT4 vector backbone) containing CD28 and CD3ζ genes (tMUC1-CAR). To test specificity of the tMUC1-CAR, we generated a control CAR (CTL-CAR), in which TAB004 scFv sequence is removed. T cells expressing CTL CAR construct is referred to as CTL T. Furthermore, to visualize surface expression of CAR constructs on T cells, we generated mKate fluorescent-tagged CARs named CAR-mKate and CTL-mKate, in which mKate2 gene is fused to the C terminus of CD3ζ of tMUC1-CAR and CTL-CAR respectively. We also used uninfected T cells (designated as mock T cells) as another control. The representative dot plot graphs show ~42% myc tag positive cells in both CD4+ and CD8+ human primary T cells by day 12 after infection (figure 1B). On average, 42% of T cells expressed tMUC1-CAR. CAR surface expression on T cells was visualized using DeltaVision microscopy. Bright field and florescent images of the entire population of CAR-mKate expressing cells are shown in figure 1C (top panel). The projection image (bottom left), and a single z stack image (bottom right) of the CAR T cell is shown in figure 1C (bottom panel). Cell nuclei were stained blue with live cell stain Hoechst. A distinct red ring indicates CAR expression on the cell surface and confirms even distribution of CAR across the cell membrane, with no significant irregular patch or co-localization (figure 1C bottom right).

**Figure 1.**
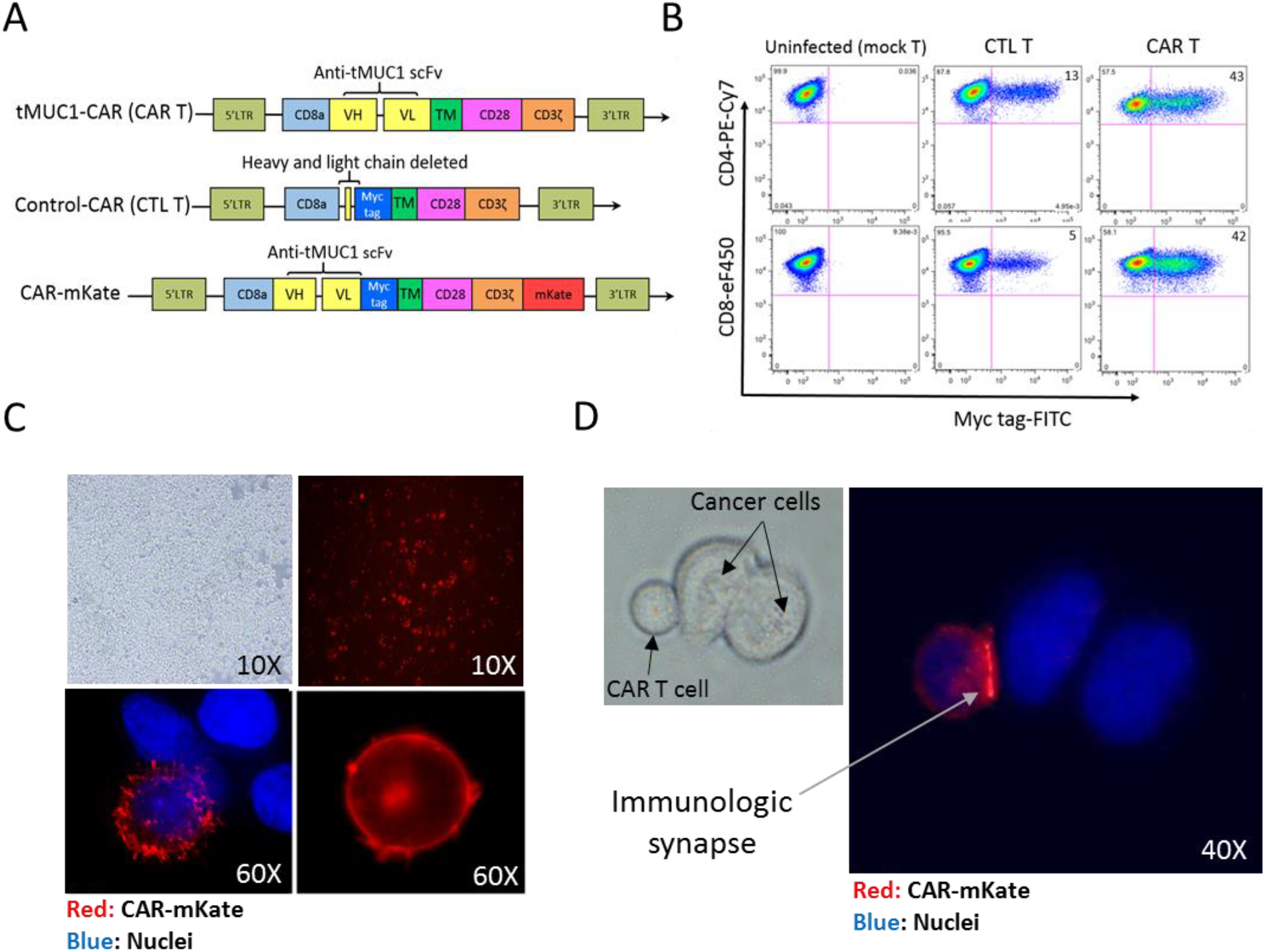
CAR architecture, expression on engineered T cells and binding of CAR-T cells to target cells. **A**. The architecture of three different CAR constructs used in this study. In the original construct (tMUC1-CAR), scFv of TAB004 antibody is linked to CD28 transmembrane (TM) domain followed by CD28 and CD3ζ intracellular domains in a retroviral plasmid. CD8a leader sequence was used as signal peptide for cell membrane expression of the CAR. In the CTL-CAR construct, scFv of TAB is removed. In the CAR-mKate construct, mKate2 gene was fused to the C-terminus of CAR flanking with a GA linker. **B.** CTL-CAR and tMUC1-CAR expression were measured by flowcytometry using FITC-conjugated anti-myc tag antibody, and expression level in CD4+ and CD8+ primary T cells on day 12 after infection are shown. On average 42% of T cells expressed tMUC1-CAR. **C**. Bright field (top left) and fluorescent image (top right) of live T cells expressing CAR-mKate plated in 35mm poly-D-lysine coated MatTek dish imaged by DeltaVision workstation (Applied Precision). Projection image of a T cell expressing CAR-mKate (bottom left), and one Z image of the CAR-mKate T cell (bottom right) illustrating the ring-like structure around the cells formed by CAR-mkate expression, which indicates even distribution of CAR molecules on the T cell membrane. **D.** Light and fluorescent image of CAR-mKate T cells binding to MUC1 expressing cancer cell (HPAFII). HPAFII cells were incubated with CAR-mKate T cells for 4 hours, then T cell were remove, HPAFII cell was washed and imaged using DeltaVision microscope. The intense red signal observed between CAR T cell and HPAFII indicates co-localization and strong binding of CAR molecules which suggests forming an immunological synapse. Nuclei are stained with Hoechst nuclei blue dye in C and D.

We have previously shown that TAB004 Ab can detect tMUC1 in >85% of malignant PDA tissues [38]. To test binding of tMUC1-CAR T cells to PDA cells, CAR-mKate engineered T cells were co-cultured with MUC1 expressing PDA cell line (HPAFII) for 4 hrs and imaged using DeltaVision microscope. Strong binding to the target HPAFII cells is observed (figure 1D); however, when CTL-mKate engineered T cells were co-cultured with the same target cells, no binding to target cells was observed (data not shown). An intensified red signal was observed where CAR T cell binds HPAFII cells, verifying co-localization of CAR molecules at the site of contact. This localization suggests formation of an immunologic synapse between the CAR T cell and target cell (figure 1D).

### tMUC1 CAR T cells show robust cytotoxicity against PDA cells but not normal cells

A panel of human PDA cell lines were used in this experiment that expressed varying levels of MUC1. BxPC3 cells overexpressing full-length MUC1 (BxPC3-MUC1) or vector alone (BxPC3-Neo) were included to further determine if CAR T cell cytotoxicity was dependent on antigen expression. Expression level of MUC1 gene and protein was assessed using RT-PCR (figure 2A) and flowcytometry (figure 2B) respectively. Cells were categorized into three groups of low MUC1, moderate to high MUC1 and high MUC1 according to flowcytometry data (figure 2B). According to RT-PCR and flowcytometry data, most PDA cell lines express some level of MUC1. Jurkat cell line was used as negative control for MUC1 gene expression in RT-PCR. Human pancreatic normal epithelial cell line, HPDE (H6c7) was used as negative control in flowcytometry and cytotoxic assays. CTL T and uninfected T cells showed similar cytotoxic activity against target cells (figure S1A); therefore, uninfected T cells (designated mock T) were used for all cytotoxic assays. PDA cell lines were incubated with CAR T or mock T cells for 24, 48 and 72 hrs at different Target: Effector (T:E) ratios and survival of target cells was measured using MTT assay. Percent survival was normalized to mock T cells and calculated as 100-{(mock T OD – CAR T OD)/mock T OD ×100}. Figure 2C shows the percentage of surviving target cells 72 hrs post T cell treatment at T:E ratio of 1:10. Majority of the tested PDA cells were efficiently killed by CAR T cells especially the high MUC1 and moderate to high tMUC1-expressing cells (80-95% killing) except for HPAFII and CFPAC (~20% and 50% killing). Interestingly, one of the low tMUC1-expressing cell line, Capan-1 also responded well to CAR T cell cytolysis. The normal pancreatic cell line, HPDE, showed 100% survival post co-culture with CAR T cells.

**Figure 2.**
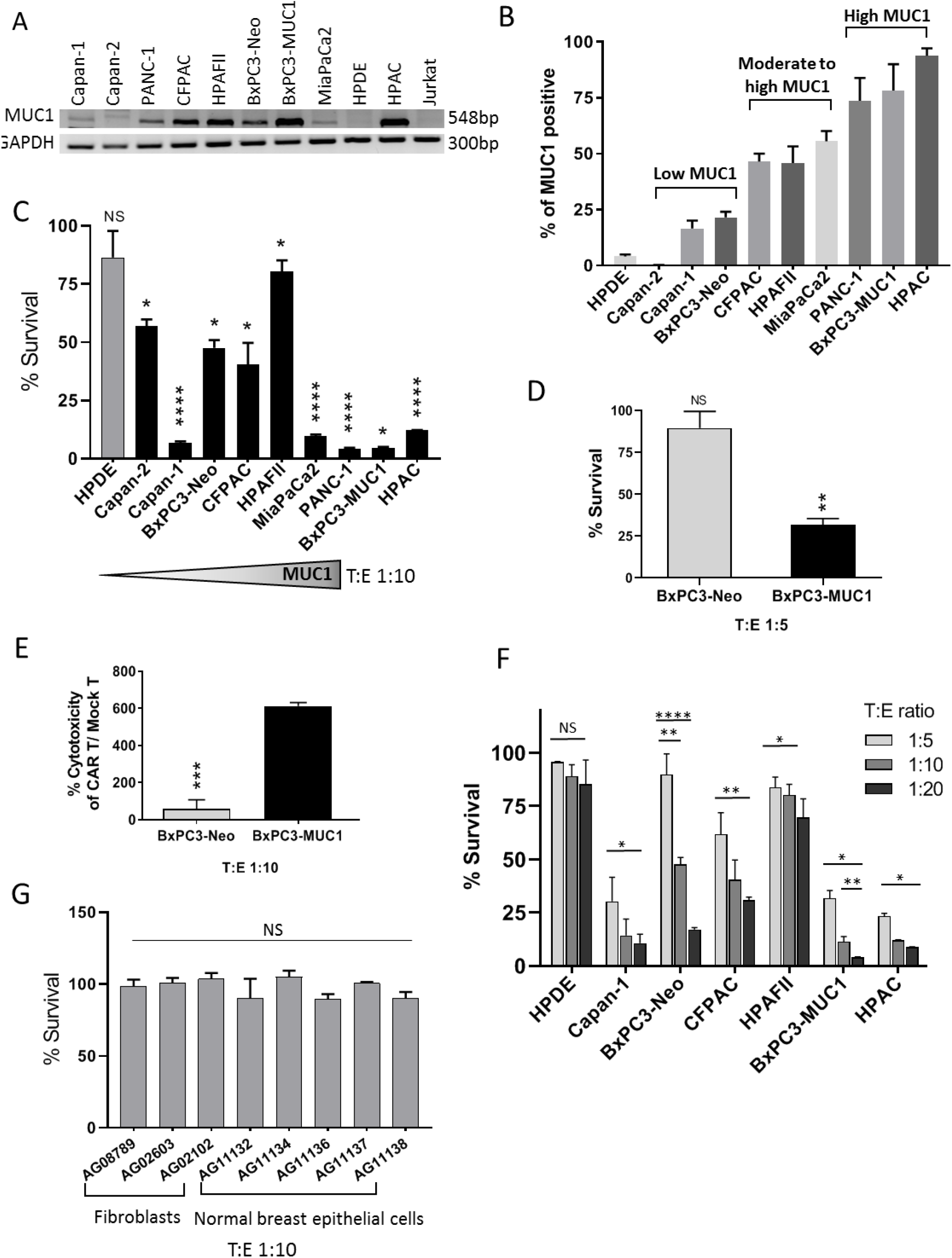
tMUC1-CAR T cells show robust cytotoxicity against PDA cells but not normal cells. **A.** The mRNA level of human MUC1 in a panel of PDA cell lines acquired by RT-PCR. Jurkat cells mRNA was used as negative control. **B.** Surface MUC1 expression in a panel of PDA cell lines detected by TAB004 Ab staining. Cancer cells were categorized into 3 groups according to their MUC1 level. **C.** The percentage survival of 9 PDA cell lines when treated with CAR T cells measured by MTT assay. Percentage survival of cancer cells treated with CAR T cell was normalized to the mock T cell (uninfected). Cancer cells are ordered from low to high MUC1 (left to right). HPDE cell was used as normal control cell line. All PDA cells show significant reduction in survival after treated with CAR T cells. T:E ratio of 1:10 and 72 hrs incubation was applied to all cell lines. Student’s t-test. **D.** The percentage survival of BxPC3-Neo and BxPC3-MUC1 treated with CAR T cells for 72 hrs at T:E ratio of 1:5. BxPC3-Neo stays intact when treated with low dose of CAR T cells (T:E 1:5), while BxPC3-MUC1 is effectively killed. Student’s t-test, ** P=0.0016. **E.** Spontaneous killing of BxPC3-MUC1 cells by CAR T cells within 24hours measured by an LDH-based technique, Cytotox assay. CAR T cells show significantly higher level of cytotoxicity against BxPC3-MUC1 cells compared to BxPC3-Neo cells. Student’s t-test, *** P=0.0004. **F.** The percentage survival of PDA cells and normal pancreatic epithelial cell line (HPDE) when treated with different dose of CAR T cells. Data shows that CAR T cell killing is dose dependent. By increasing the dose of CAR T cells, more killing is observed in PDA cells, while the survival of normal cell (HPDE) even at T:E of 1:20 stays unchanged. Two-way ANOVA-Multiple comparisons. **G.** The percentage survival of a panel of human normal primary cells including fibroblasts and breast epithelial cells obtained from healthy donors treated with CAR T cells for 72 hrs at T:E 1:10. There is no significant reduction in the survival level of normal primary cells when treated by CAR T cells. All of the survival levels were normalized to mock T cells. Student’s t-test. Error bars, SEM. * p< 0.05, ** p< 0.01, *** p< 0.001, **** p< 0.0001.

The sensitivity of the PDA cells to CAR T cell killing cannot be accurately compared between the different cell lines, since each cell line has distinct genetic makeup, which endows them different intrinsic resistance levels. To investigate this, we used BxPC3-MUC1 and BxPC3-Neo cells. These two cell lines are identical in all aspects except for their MUC1 level. As shown in figures 2C, D, and E, BxPC3-MUC1 cells are significantly more sensitive to CAR T cell killing as compared to BxPC3-Neo cells. At a T:E ratio of 1:10 (figure 2C), ~100% of BxPC3-MUC1 cells are killed by CAR T cell treatment while ~40% of BxPC3-Neo cells are killed. This 40% killing with a high T:E ratio (1:10) in BxPC3-Neo cells was possibly because of the existing endogenous MUC1 expression in these cells (figure 2A, B). However, the killing effect was completely negated when T:E ratio was lowered to 1:5 (figure 2D). The cell survival data was further confirmed using a different cell cytoxicity assay (CytoTox 96^®^ Non-Radioactive Cytotoxicity Assay). Percent cytotoxicity of CAR T cells/Mock T cells show significant killing of BxPC3-MUC1 but not BxPC3-Neo cells (figure 2E). Taken together, results clearly suggest a critical correlation between antigen expression levels and efficacy of CAR T cells. Next, we evaluated whether the efficacy of CAR T cell killing is dose dependent. Indeed, increasing ratio of CAR T cells to target PDA cells resulted in dose dependent killing of the target cells (figure 2F). CAR T cells had no effect on normal pancreatic epithelia cells (HPDE) at 1:5, 1:10 or 1:20 T:E ratio while the same CAR T cell effectively killed the high-MUC1 expressing PDA cell lines, at all T:E ratios (figure 2F).

Lastly, we tested if tMUC1-CAR T cells kill other normal epithelial cells or fibroblasts. We performed the same cell survival assay using eight different primary cells as targets (granted from Coriell institute) at a 1:10 T:E ratio and 72 hrs incubation with CAR T cells. Three fibroblasts from three different tissue origins and 5 breast epithelial cells (from mammoplasty) derived from different healthy donors were tested. All normal cells showed 100% survival when exposed to CAR T cells for 72 hrs (figure 2G). These data suggest that the tMUC1-CAR T cells are non-toxic toward normal cells while robustly killing most PDA cell lines. This is of utmost importance since CAR T cell toxicity in patients with epithelial tumors has raised major concerns.

Interaction of CAR T cells with BxPC3-MUC1 and BxPC3-Neo cells were recorded overnight using GE DeltaVision OMX-SR microscope and the video was created using the ImageJ program. BxPC3-MUC1 cells were attacked and killed by CAR T cells in 7/9 spots, while only 2/9 spots showed killing in BxPC3-Neo plate. Red staining is indicative of dead cells that have absorbed PI dye present in the media. Videos are available as supplementary data: video.

### tMUC1-CAR T cells produce IFN-γ and granzyme B upon activation and antigen recognition

To determine the mechanism of T cell activation and function, T cells were co-cultured with target cells for 72 hrs and supernatants were tested for IFN-γ and granzyme B production by specific ELISAs (figure 3A, B). In addition, intracellular production of IFN-γ by T cells after co-culture was measured by flowcytometry (figure S1B). Results show that even before exposure to cancer cells, activated T cells produced some level of IFN-γ cytokine (intracellular and released); however, when exposed to cancer cells, IFN-γ secretion by CAR T cells significantly increased. As may be expected, ELISA data showed that higher MUC1 expressing PDA cells triggered higher levels of IFN-γ and granzyme B release by CAR T cells. CTL T and mock T cells released negligible amount of IFN-γ and granzyme B into the media even after exposure to target cells. Other controls including supernatants from a) cancer cells alone and b) Jurkat T cells, as well as c) media alone, showed undetectable amount of released IFN-γ and granzyme B by ELISA (figure 3A, B). Intracellular level of IFNγ (figure S1B) and granzyme B (data not shown) in CAR T cells post exposure to PDA cells showed similar results as the ELISA. We further tested levels of intracellular perforin in CAR T cells before and after exposure to target cells by flowcytometry and the results showed no difference between resistant and sensitive target cells (figure S1C).

**Figure 3.**
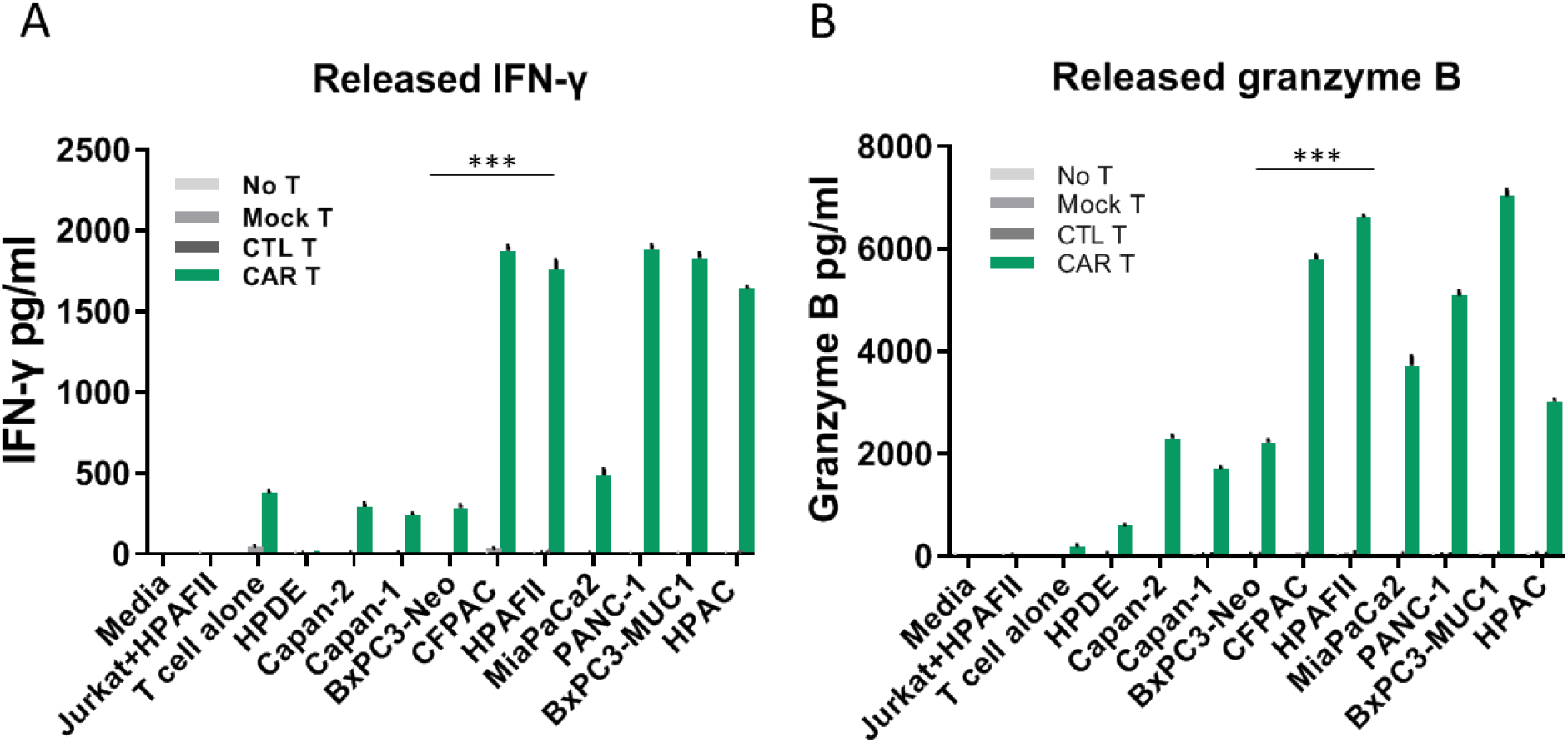
tMUC1-CAR-T cells produce IFN-γ and granzyme B upon activation and antigen recognition. The amount of released IFN-γ **(A)** and granzyme B **(B)** in the co-culture media of CAR T cells and cancer cells measured by sandwich ELISA. Controls include supernatant of 1) cancer cells alone, 2) Jurkat and HPAFII cells co-culture, 3) T cells alone as well as 4) media alone. The cancer cells are ordered based on their MUC1 level from left to right (low to high MUC1). tMUC1-CAR T cells exposed to cancer cells produce significant amount of IFN-γ and granzyme B, while CTL T and mock T cells exposed to cancer cells produce negligible amount of IFN-y and granzyme B. CAR T cells exposed to normal epithelial cell line, HPDE, did not produce noticeable amount of IFN-γ and granzyme B. Error bars, SEM. Student’s t-test, P value < 0.0005 for CAR T cells vs. mock T.

### tMUC1-CAR T cells control pancreatic tumor growth *in vivo*

To investigate the efficacy of CAR T cells in hampering tumor growth *in vivo*, orthotopic mouse model of human PDA was stablished by injecting 0.5×10^6^ MiaPaCa2-Luc cells into the pancreas of NSG mice. Seven days post-surgery, mice were imaged using IVIS and tumor presence was confirmed. On day 8 post-surgery, mice were randomized into two groups based on their baseline luminescence intensity. One group received IV injection of Mock T and the other received CAR T cells (10×10^6^ per mouse) (figure 4A). Tumor growth was monitored using weekly IVIS imaging (data not shown). On day 68 post-surgery, mice were sacrificed and tumors were harvested. Mock T cell-treated mice had significantly larger tumors than CAR T cell-treated mice (figure 4B). Many metastasis lesions were present on organs in the abdominal cavity of mock T cell-treated mice, whereas CAR T cell-treated mice had more confined tumors (data not shown). Tumor wet weights were measured and results showed significant difference between the CAR T and mock T cell treated groups (P value=0.0476, figure 4B).

**Figure 4.**
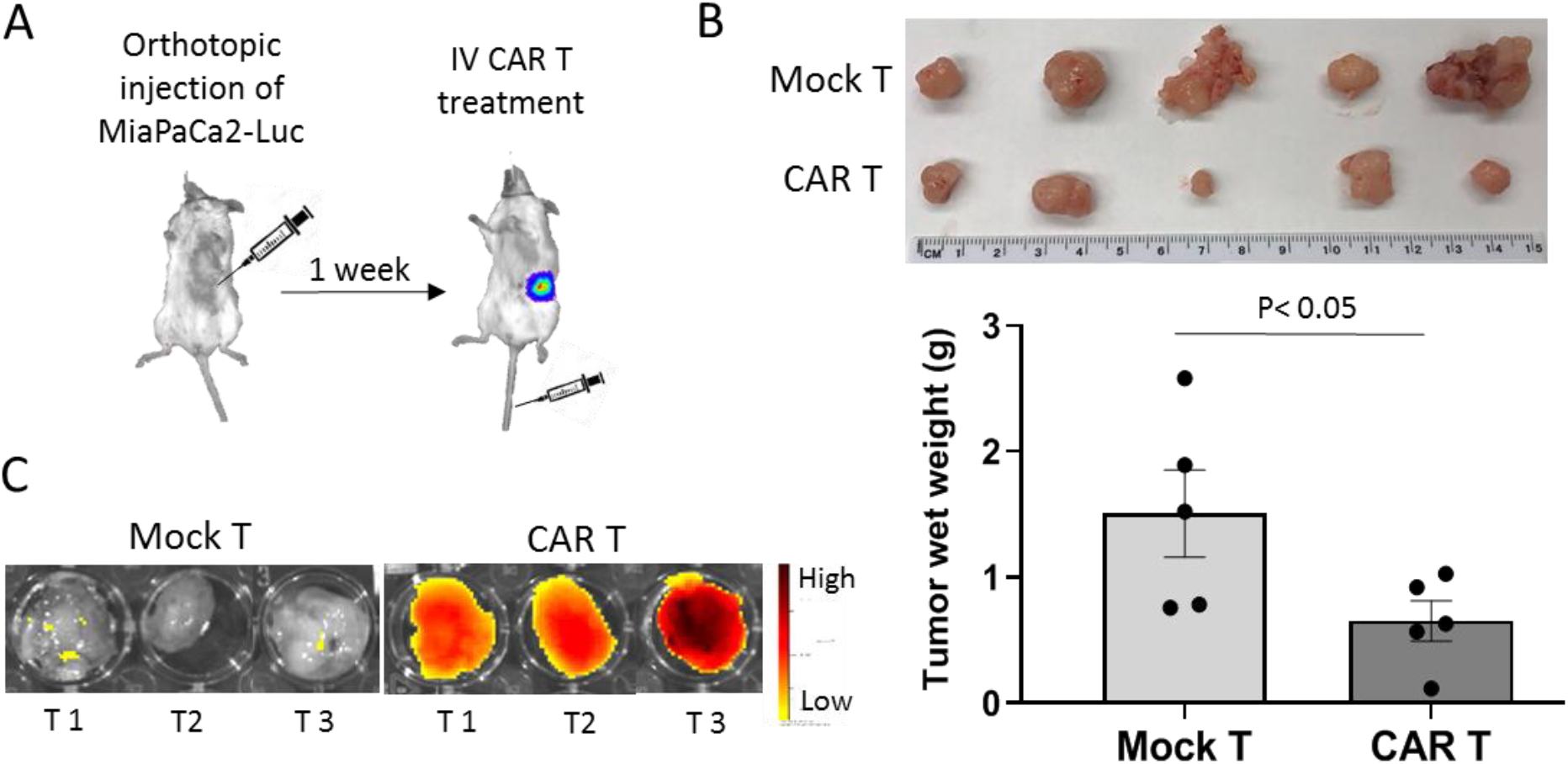
tMUC1-CAR T cells control pancreatic tumor growth *in vivo.* **A**. Stablishing the mouse model of human PDA using orthotopic injection of MiaPaCa2-Luc cancer cells into the pancreas. Using aseptic techniques, a small incision was made in abdominal area off midline just above the pancreas of the mice. Pancreas was gently retracted and injected with 0.5×106 MiaPaCa2-Luc (luciferase positive) tumor cells. 7 days post-surgery, tumor presence was confirmed using IVIS imaging. On day 8, mice were randomized into two groups and injected IV with 10×106 mock or CAR T cells. Images were taken weekly at 8 minute after luciferin injection using IVIS system (data not shown). **B.** Images of the tumors harvested from mice treated with mock T or CAR T cells on day 68 post tumor inoculation (top). Tumor wet weights of the mice treated with mock T or CAR T cells on day 68 after tumor inoculation (bottom). Significance of data was evaluated using Non-parametric Mann-Whitney U test. P value=0.0476 (n=5). **C.** Visual representation of CAR T cells trafficking in the pancreatic tumors. To evaluate T cells trafficking into the fibrotic pancreatic tumor, six tumor bearing mice (day 52 post-surgery) were injected IV with either 4×106 vivotrack-680 labeled-CAR T cells or mock T cells. After 24 hrs, mice were scarified and tumors were harvested and imaged using fluorescent channel on IVIS machine with excitation=676 and emission=696 nm. The fluorescent signal acquired from tumors of mice treated with CAR T cells was significantly higher than the ones treated with mock T cells, which indicates more CAR T cells are directed to the tumor site than mock T cells. T1-3, tumor 1-3.

To see if CAR T cells can successfully traffic into the fibrotic pancreatic tumors, mock and CAR T cells were labeled with Vivotrack-680 dye and IV injected into six MiaPaCa2-Luc tumor-bearing NSG mice on day 52 post-surgery (n=3). Mice were sacrificed 24 hrs after (since Vivotrack-680 signal intensity peaks at 24 hrs), and tumors were dissected and imaged by IVIS using Vivotrack-680 channel. There was strong signal emitted from CAR T cell-injected tumors compared to the control group, which suggests that CAR T cells were able to infiltrate into and localize in the pancreatic tumor mass as early as 24 hrs after infusion, while mock T cells were not directed to the tumor mass (figure 4C).

### Deciphering the intrinsic resistance mechanism utilized by PDA cells to CAR T cell therapy: Role of IDO1 and Gal-9

To assess why some PDA cell lines are resistant to CAR T cell killing independent of their tMUC1 expression or the CAR T cell’s ability to express perforin (figure S1C), or produce IFN-γ and granzyme B (figure 3 and S1B), we considered some common immune evasion tactics used by tumor cells. We selected two highly resistant cell lines; HPAFII and CFPAC (both express similar levels of tMUC1, figure 2B). As control, we included a highly sensitive cell line, MiaPaCa2 with similar tMUC1 level.

Impairing T cells function by cancer cells is reported as a common mechanism involved in tumor immune evasion. Main factors in T cells anti-tumor cytotoxicity include IFNγ, granzyme B and perforin secretion. Results show no correlation between the amount of released IFNγ and granzyme B and levels of intracellular perforin with the resistance of tumor cells (figures 3 and S1B, C).

Another common mechanism of immune evasion is associated with tumor-induced apoptosis of effector T cells. PDL1 expressed by cancer cells can interact with PD1 receptor on T cells and trigger T cell apoptosis. To test if HPAFII and CFPAC utilize this mechanism, mock and CAR T cells were exposed to resistant (HPAFII, CFPAC) or sensitive (MiaPaCa2) cells for 24, 48 and 72 hrs and the apoptosis was assessed by Annexin V/PI staining and flowcytometry. Data shows that apoptosis of mock and CAR T cells did not significantly alter after 24, 48 and 72 hrs exposure to resistant vs. sensitive PDA cells. T cell apoptosis level at 48 hrs time-point is shown in figure 5A. Level of PD1 expression by T cells before and after exposure to resistant or sensitive cells were measured by flowcytometry and the results are shown in figure S2A, B. There was no significant difference between level of PD1 expression by mock and CAR T cells before and after co-culture with resistant or sensitive cells at 24, 48 and 72 hrs. Most PDA cells used in this study express considerable amount of PDL1; however, the PDL1 level is not correlated with the resistance of PDA cells (figure S2A, B). To confirm the results, PD1 blocking with anti-PD1 Ab was performed in combination with CAR T cell therapy on the three cell lines. We detected no improvement in killing of the resistant PDA cell lines (HPAFII, CFPAC), while sensitive cell line (MiaPaCa2) killing was enhanced (figure S3). Data suggests that PD1/PDL1 interaction may not be the major factor driving CAR T cell resistance in HPAFII and CFPAC cells.

**Figure 5.**
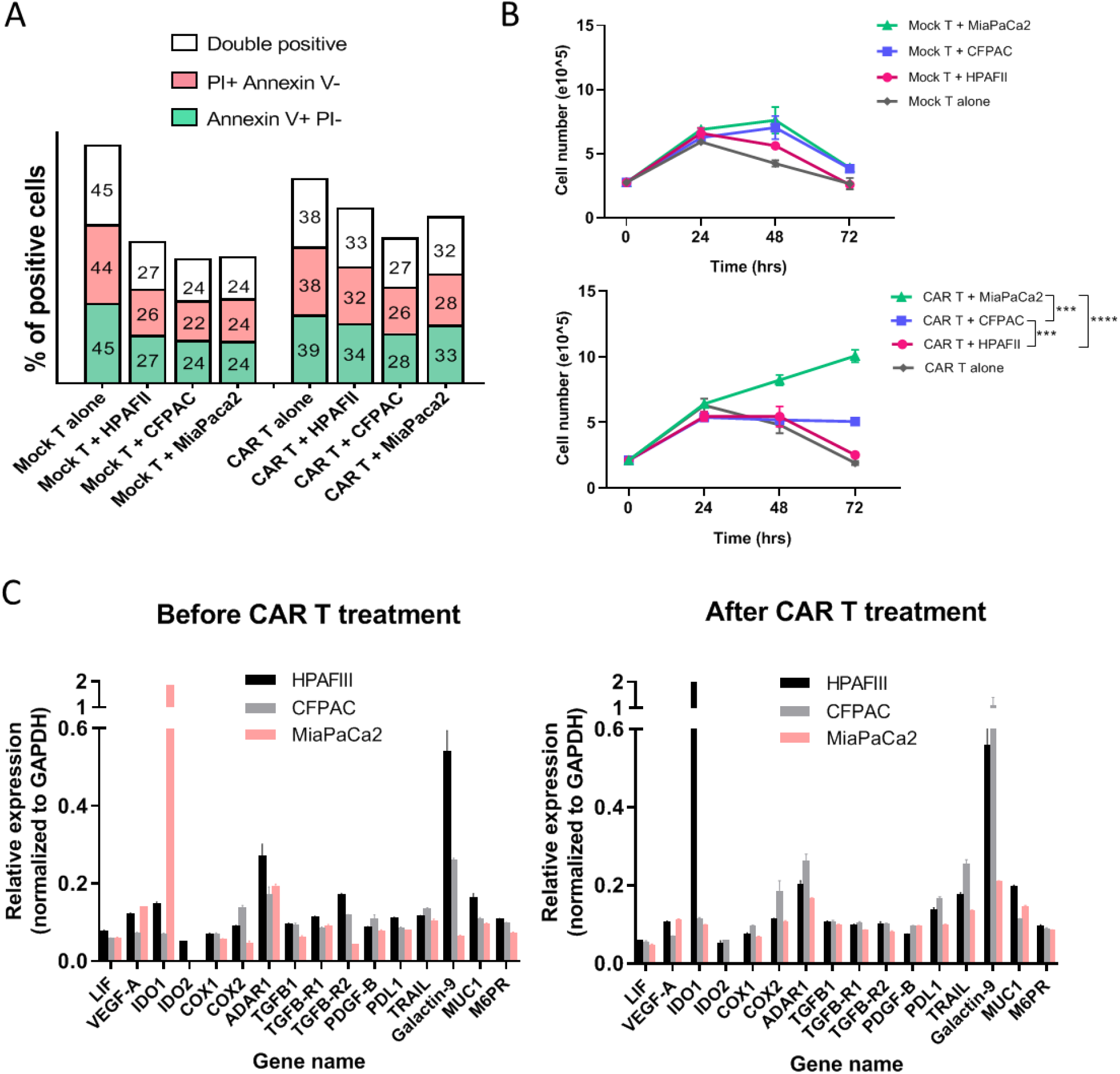
Deciphering the resistance mechanism utilized by PDA cells against CAR T cell therapy. **A.** T cells apoptosis level before and after exposure to HPAFII, CFPAC and MiaPaCa2 cells. Mock and CAR T cells were co-cultured with PDA cells and their apoptosis level was measured by Annexin V/PI staining at 24, 48 and 72 hrs post co-culture. Figure A shows percentage of positive Annexin V, PI or both in T cells after 48 hrs co-culture. There is no significant difference between apoptosis level of mock and CAR T cells when exposed to resistant vs. sensitive PDA cells. **B.** Mock and CAR T cells proliferation over time after exposure to HPAFII, CFPAC and MiaPaCa2 cells. T cells were enumerated using an automated cell counter. MiaPaca2 cells enhanced proliferation of CAR T cells after 48 and 72 hrs, whereas CFPAC and HPAFII cells hindered CAR T cells proliferation. Mock T cells didn’t show the same trend. Significance of data was evaluated using Two Way ANOVA (Multiple Comparison). Error bars, SEM.*** p< 0.001, **** p< 0.0001. **C**. q-PCR data showing relative expression level of 16 genes in HPAFII, CFPAC and MiaPaCa2 cells before (top) and after (bottom) exposure to CAR T cells. CTs are normalized to GAPDH in each sample and higher number in Y axis represents higher expression of the gene. IDOI, COX1/2, ADAR1 and galectin-9 genes expression was higher or increased after CAR T treatment in resistant PDA cells.

Because apoptosis of CAR T cells was not significantly different when co-cultured with resistant versus sensitive PDA cell lines, we enumerated cell numbers post co-culture. Proliferation of CAR T and Mock T cells was evaluated before and after co-culture with resistant (HPAFII and CFPAC) and sensitive (MiaPaCa2) PDA cells at 24, 48, and 72 hours using a cell counter. Interestingly, CAR T cell proliferation was significantly hindered when co-cultured with HPAFII and CFPAC cells by 72 hrs, while growth of CAR T cells was enhanced when co-cultured with MiaPaca2 cells. Mock T cell proliferation did not show the same trend (figure 5B).

Studies have shown that PDA cells shed MUC1 into the supernatant and that may cause impaired T cell function. It is also reported that depletion of soluble MUC1 from the tumor supernatants reversed the inhibitory effects on T cells [39]. Thus, we assessed the amount of released MUC1 by PDA cells in the co-culture media using a specific ELISA (figure S2C). Results show that only HPAC cells release high levels of MUC1, while other PDA cells shed minimal levels of MUC1. Since HPAC cell line is highly sensitive to CAR T cell treatment, we negated the role of shed MUC1 as a mechanism for antigen loss and immune escape (figure S2C).

Thus, we moved to investigate the gene expression profile of 16 genes linked to immune resistance in HPAFII, CFPAC, and MiaPaCa2 cells. Sixteen genes linked to immune resistance (based on literature) were analyzed using qPCR technique. Relative expression of each gene was normalized to GAPDH level. Figure 5C shows relative expression of each gene (normalized to GAPDH) in three cell lines before and after CAR T cell treatment. MUC1 mRNA expression in resistant vs. sensitive cells did not significantly change after CAR T cell treatment, indicating no antigen loss through gene downregulation, and therefore may not account for the immune evasion in the HPAFII and CFPAC PDA cells (figure 5C). Most of the genes were expressed at low levels in resistant cells except indoleamine 2, 3-dioxygenases-1 (IDO1), Cyclooxygenase 1 and 2 (COX1/2), Adenosine deaminases acting on RNA (ADAR1) and Gal-9. According to qPCR data, IDO1 gene expression in HPAFII cells was significantly increased (69-fold increase) after treatment with CAR T cells, while its level declined in MiaPaCa2 cells. COX1 and 2 expressions were slightly higher in HPAFII and CFPAC, and their level increased after CAR T treatment. The expression of ADAR1 gene was higher in HPAFII cells compared to MiaPaCa2 before treatment, and expression of this gene was only increased in CFPAC after CAR T treatment. Gal-9 gene expression was also high in both resistant cells before and after CAR T cell therapy. These genes are appropriate candidates to further study as some of the important players in immune resistance of HPAFII and CFPAC cells. Hence, we combined CAR T cell treatment with inhibitors or blocking antibodies to the above-mentioned molecules.

## Battling the resistance of PDA cells with combination therapy

### Targeting resistance related genes with small molecule inhibitors and blocking antibody

IDO1 was one of the candidate genes involved in immune resistance. IDO1 function can be inhibited by 1-Methyl-D-tryptophan (1-MT) drug. Three cell lines were treated with 1-MT drug at three different concentrations for 1 day followed by 3 days of mock or CAR T cell treatment. CAR T cells plus drug killing was normalized to mock T cell plus drug, and the asterisk shows significant difference between the CAR T cells plus drug (combination therapy) and CAR T cells alone. Results showed a significant reduction in survival of HPAFII and CFPAC cells by combination of CAR T cells and 1-MT therapy in a dose dependent manner compared to CAR T alone or 1-MT alone therapy, while MiaPaCa2 cells showed no significant difference in survival (figure 6). Data suggests that IDO1 may be one of the major factors causing immune resistance in HPAFII and CFPAC cells and targeting IDO1 along with CAR T cell therapy may enhance the treatment efficacy in resistant PDA cells.

**Figure 6.**
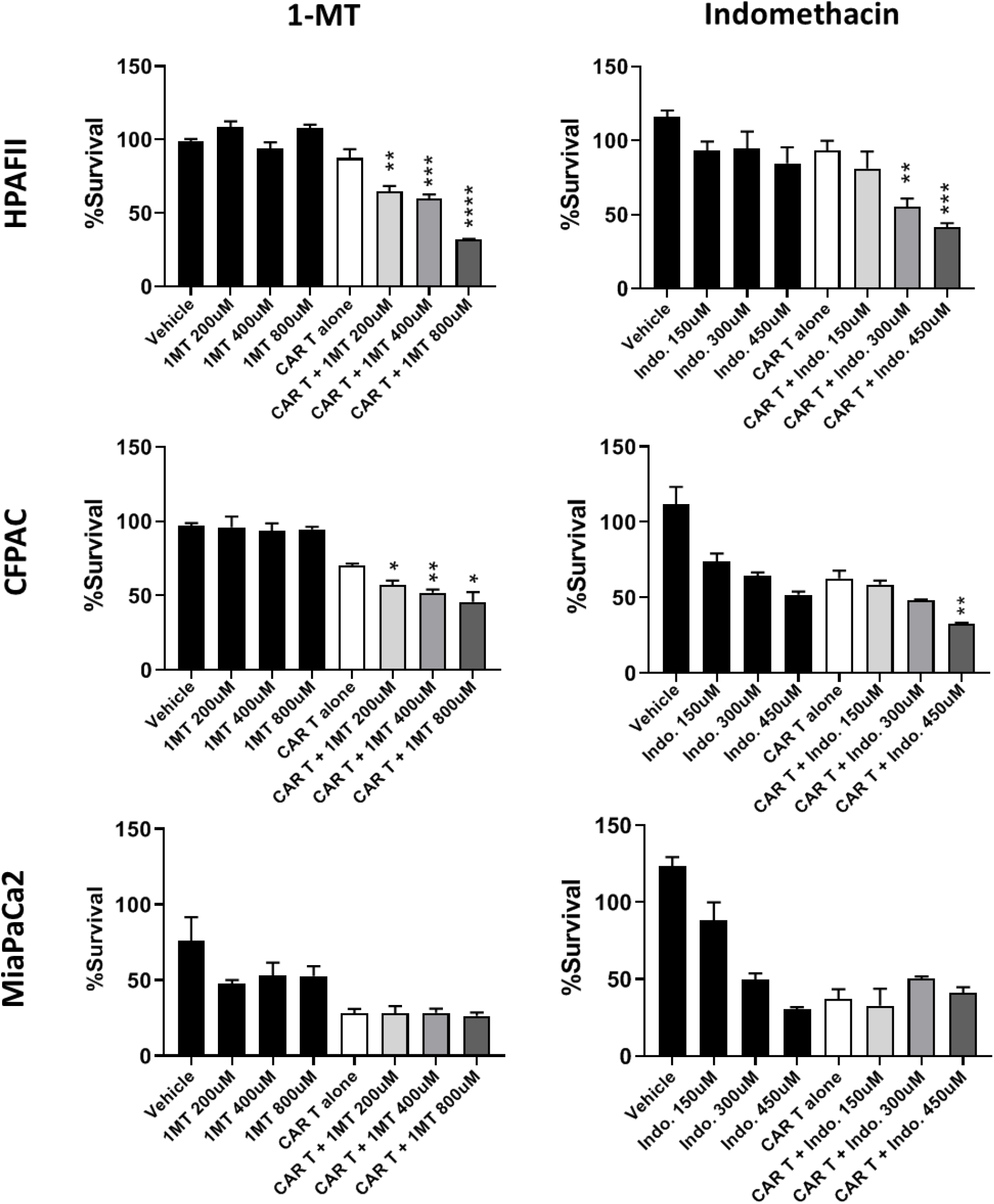
Targeting resistance related genes with small molecule inhibitors. HPAFII, CFPAC and MiaPaCa2 cells were pre-treated with IDO inhibitor (1-MT) or COX1/2 inhibitor (indomethacin) for 24 hrs, then drugs were removed and PDA cells were co-cultured with mock T or CAR T cells for 72 hrs at T:E ratio of 1:10. Percentage survival was measured using MTT assay and normalized to mock T. HPAFII and CFPAC killing by CAR T cells was significantly improved when pre-treated with 1-MT and indomethacin; while MiaPaCa2 cell did not respond to the combinational treatment. Student’s t-test comparing CAR T + drug group to CAR T alone group. * p< 0.05, ** p< 0.01, ***, p< 0.001, **** P< 0.0001.

Several studies have shown the importance of COX1 and COX2 in causing resistance of cancer cells to immune therapy [40] [41]. Therefore, celecoxib (specific COX-2 inhibitor) and indomethacin (COX1 and 2 inhibitor) were used in combination with CAR T cells. Both HPAFII and CFPAC cells showed reduction in survival when treated with indomethacin and CAR T cells, while MiaPaCa2 cells showed no difference in survival (figure 6). Celecoxib did not change the efficacy of CAR T cells (Data not shown).

Next candidate gene for resistance was ADAR1. qPCR data showed an elevation in ADAR1 gene expression level in resistant cells compared to MiaPaCa2. PDA cells were treated with EHNA drug (ADAR1 inhibitor) for 24 hrs before adding CAR T cells, and cell survival was measured at 72 hrs post co-culture. Combination therapy with EHNA drug did not result in significant reduction in the survival of the target cells (data is not shown). Hence, we inferred that ADAR1 may not play an important role in immune resistance of HPAFII and CFPAC cells.

Gal-9 was another gene that showed increased expression in the resistant cells. Gal-9 is a tandem-repeat galectin interacting with Tim3 receptor on T cells [42]. To neutralize the effects of Gal-9, a blocking Ab, 9M1-3 (BioLegend) was used in combination with CAR T cells and the results are shown in figure 7. HPAFII and CFPAC cells are efficiently targeted by CAR T cells when Gal-9 checkpoint inhibitor is added to the co-culture media compared to CAR T cell or 9M1-3 treatment alone. In contrast, survival of MiaPaCa2 cells was not affected by the combination. Data suggests Gal-9 –Tim3 interaction may play an important role in immune evasion by resistant PDA cells.

**Figure 7.**
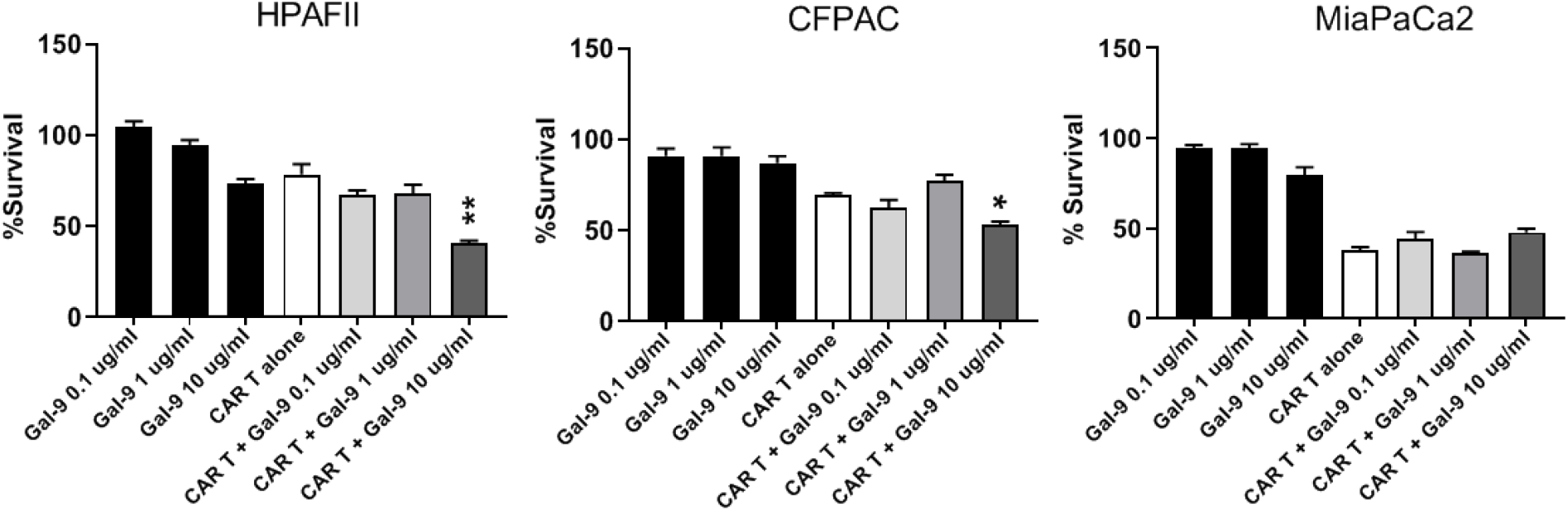
Targeting resistance related genes with anti-Gal-9 blocking antibody. Percentage survival of HPAFII, CFPAC and MiaPaCa2 cells after treatment with CAR T alone, Gal-9 blocking antibody alone, and combination of CAR T cell and Gal-9 blocking Ab. Anti-Gal-9 Ab was added at 3 different concentrations to the co-culture media of PDA cells and mock or CAR T cells (T:E 1:10). Percentage survival was obtained using MTT assay and data was normalized to mock T. HPAFII and CFPAC survival were reduced with combination of CAR T and anti-Gal-9 blocking Ab; while MiaPaCa2 did not respond to the combination therapy. Student’s t-test comparing CAR T + anti-Gal-9 Ab group to CAR T alone group. * p< 0.05, ** p< 0.01.

### tMUC1-CAR T cells work synergistically with common chemotherapy drugs to kill resistant PDA cells

Another approach to break resistance of target cells to CAR T cells is pre-sensitizing them with common chemotherapy drugs. Three widely used drugs for PDA include gemcitabine (GEM), 5-Fluorouracil (5FU) and paclitaxel (PTX). GEM is analog of deoxy-cytidine, inhibits DNA synthesis, PTX suppresses microtubule detachments from centrosome, and 5-FU inhibits Thymidine synthase. We treated PDA cells with the drugs for 24 hrs prior to CAR T cell treatment. After 72 hrs of CAR T treatment, target cell viability was measured using MTT assay. Results demonstrate significantly enhanced sensitivity of HPAFII cells to CAR T cell treatment when pre-exposed to low-dose chemotherapy drugs (figure 8, top row). All three drugs improved killing of HPAFII cells with CAR T cells, while only 5-FU showed the same effect on CFPAC cells. GEM and PTX did not assist the CFPAC sensitivity to CAR T cells (figure 8, bottom row). Data suggests that pre-sensitizing the resistant PDA cells with low dose of appropriate chemotherapeutic drug may enhance the efficacy of CAR T cell treatment.

**Figure 8.**
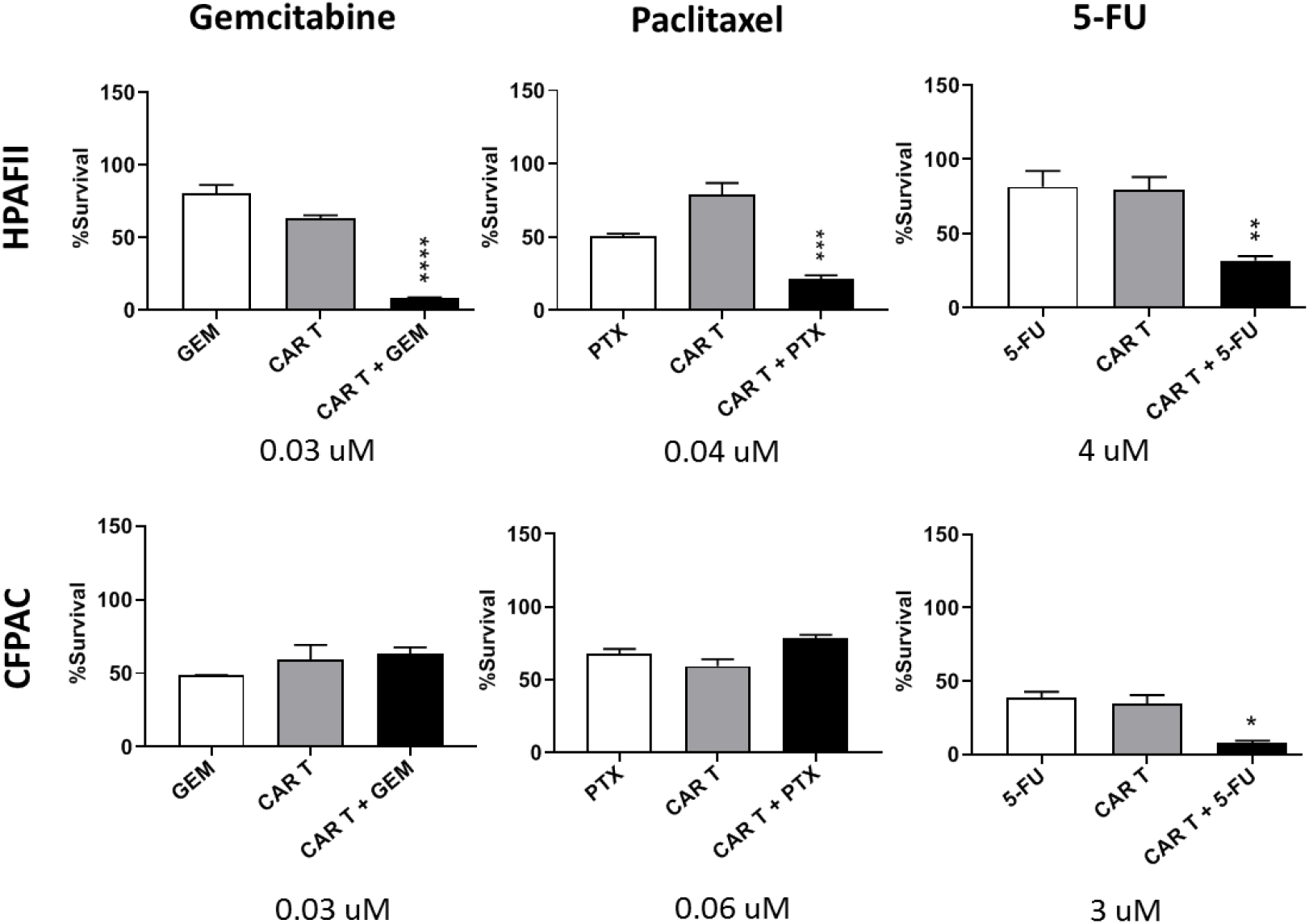
tMUC1-CAR T cells work synergistically with common chemotherapy drugs to kill resistant PDA cells. Percentage survival of two resistant PDA cells, HPAFII and CFPAC, treated with combination of CAR T and chemo drugs. HPAFII and CFPAC were exposed to chemotherapy drugs (gemcitabine, paclitaxel or 5-FU) for 24 hrs at indicated concentrations, then co-cultured with mock or CAR T cells at T:E ratio of 1:10. Survival level was measured using MTT assay and data was normalized to mock T. Student’s t-test comparing CAR T + drug to CAR T alone group, ** P> 0.0021, *** P<0.0004, **** P < 0.0001

## Discussion

We present preclinical data showing efficacy of a novel TAB004-derived CAR T cell targeting tMUC1 against a panel of human PDA cell lines. Data demonstrates that some PDA cells remain highly resistant to CAR T cell therapy. Thus, we determined the genes that may be involved in the immune resistance. Data clearly implicate IDO1, Gal-9 and to a lesser extent, COX genes as mechanisms of CAR T cell resistance. To our knowledge, this is the first study to show that blocking the function of IDO1 or Gal-9 can enhance CAR T cells therapy against PDA cells. We also report that pre-treating the resistant HPAFII cells with standard drugs such as PTX, GEM, and 5FU enhanced CAR T cell efficacy. However, for CFPAC cells, only pre-treatment with 5FU enhanced CAR T cell efficacy. Data suggest that not all PDA cells are alike and biomarkers such as IDO1 and Gal-9 may be useful tools to determine response to CAR T cell therapy.

Recently, a MUC1 CAR T cell using the scFv from a different MUC1 Ab was shown to be efficacious against breast and pancreatic tumors [21] [24]. Our data corroborates that tumor associated MUC1 remains a critical targetable antigen in PDA. This study differs from previously reported data in 3 major ways. 1) use of variable fragment from a highly specific tMUC1 Ab, TAB004, that does not recognize normal MUC1; 2) testing the efficacy of the tMUC1-CAR T cell against a large panel of PDA cell lines and normal epithelial and fibroblastic cells, and 3) deciphering the potential intrinsic immune tolerance mechanisms utilized by PDA cells to resist CAR T cell therapy. Furthermore, this is the first study indicating the role of Gal-9 in conferring resistance in PDA cells to CAR T cell treatment and that ADAR1 and COX pathway were less critical in conferring resistance.

CAR T cell therapy, though extremely successful in treating hematopoietic cancers, has not gained momentum in the treatment of solid tumors [9]. This is primarily because of limited selection of tumor specific antigen and a dearth of specific antibodies against them. CAR based on antibodies which recognize a shared tumor antigen such as ERBB2, have led to lethal outcomes [34]. Other MUC1 antibodies may be limited by their specificity, wherein they bind to tumor and normal MUC1. TAB004 has high specificity and binding to tMUC1 [29] and spares binding to normal MUC1. We present data that clearly shows that the tMUC1-CAR T cells do not kill normal cells but effectively kill majority of tMUC1 expressing PDA cells. Recently, our group has published the effectiveness of tMUC1-CAR T cell efficacy against TNBC cells *in vitro* and *in vivo* [43]. This highlights the potency of tMUC1-CAR T cells as a potential therapy for multiple adenocarcinomas.

Despite many PDA cells being efficiently destroyed by CAR T cells, few PDA lines were highly resistant. This was not surprising. Human PDA is known to be immunologically cold and refractory to the treatments. Thus, deciphering mechanisms involved in PDA immune-resistance is of cardinal importance. Majority of combination therapy to date have focused on targeting the PD1/PDL1 axis using blocking antibodies [44], but in our study with HPAFII and CFPAC cells, we saw no evidence supporting enhanced CAR T cell efficacy when combined with anti-PD1 antibody [45]. A study by Koyama et al [46] has shown that failure of PD1 monotherapy blockade or PD1 adaptive resistance in lung adenocarcinoma is associated with upregulation of alternative immune checkpoint molecules particularly Tim-3. This may explain why PD1 blockade in our model did not improve HPAFII and CFPAC killing, dissimilar to MiaPaCa2 cells. Tim-3 upregulation in HPAFII and CFPAC cells with elevated levels of its ligand, Gal-9, may potentially deteriorate CAR T cells function.

Multiple factors can dictate the resistance of a tumor cells to a treatment. High-throughput screening is required and yet it may result in inconclusive data. In this study, we focused on key proteins and genes known to be associated with immune tolerance. Sixteen genes were analyzed using qPCR. Among those, IDO1, an enzyme that catabolizes tryptophan to kynurenine acid thereby provides metabolic advantage to cancer cells against T cells, was found to be of utmost importance [47, 48]. IDO1 contributes to peripheral immune tolerance and evasion of tumors by downregulating T cell metabolism. Several studies have confirmed that IDO1 and the downstream tryptophan catabolites inhibit T cell proliferation, thereby suppressing T cell function [49–52]. As shown in our study, CAR T cell proliferation was stunted when co-cultured with resistant PDA cell line (figure 5B). This reduction in proliferation could be due to elevated IDO1 activity in cancer cells and the release of tryptophan catabolites in the co-culture media. Accordingly, neutralizing the effect of IDOI by 1-MT drug resulted in improved lysis of HPAFII and CFPAC cells by CAR T cells.

Another gene with distinct overexpression in resistant cells was Gal-9. Gal-9 is a tandem-repeat type galectin that like other galectins modulate multiple biological functions such as cell adhesion and aggregation. Although the role of Gal-9 in immunity is controversial [53], many studies confirmed that Gal-9 negatively regulates T cells via interaction with Tim-3 receptor [42, 54, 55]. Tim-3 is a negative regulatory immune checkpoint expressed on T cell which is known to inhibit the immune responses of TH1 cells and plays an important role in immune exhaustion of T cells [56]. Studies have shown that interaction between Gal-9 and Tim-3 triggers cell death in effector Th1 cells [57] and in Tim-3^+^CD8^+^ Tumor infiltrating lymphocytes [58]. However, not all Gal-9-Tim-3 interactions result in cell death, as Gal-9 was found to increase Tim-3-mediated IFN-γ production in an NK cell line [59]. We have evaluated the level of Tim-3 expression on CAR T cells when exposed to resistant and sensitive target cells, but no significant difference was detected (data not shown). However, we found that targeting Gal-9 with a blocking antibody reduced tolerance of resistant HPAFII and CFPAC to tMUC1-CAR T cell therapy. Hence, Gal-9 immunosuppressive role may be mediated through other less known mechanisms.

ADAR1 was noticeably overexpressed in HPAFII and CFPAC cells. ADAR1 regulates the biogenesis of members of the miR-222 family and thereby ICAM1 expression, which ultimately leads to immune resistance [60]. Loss of function of ADAR1 in tumor cells strongly sensitizes tumors to immunotherapy and overcomes resistance to PD1 checkpoint blockade [61]. Surprisingly, in this study, blocking ADAR1 function with EHNA drug did not result in breaking resistance of HPAFII and CFPAC cells to CAR T cells treatment.

Finally, COX1/2 are enzymes catalyzing the synthesis of prostaglandin E2 (PGE2), a major player in inflammation, angiogenesis, and immunosuppression in cancer [62, 63]. COX2 is often overexpressed in cancer cells and is associated with progressive tumor growth, as well as resistance of cancer cells to conventional chemotherapy, immunotherapy, and radiotherapy [62]. COX1/2 inhibitors have been used in combination with anti-cancer agents and immunotherapy against cancers [64]. COX1 and COX2 were slightly higher in the resistant vs. sensitive PDA cells. Indeed, indomethacin significantly improved CAR T cell efficacy when HPAFII cells were pre-treated with the drug. To our surprise, celecoxib had no such effect. This may indicate COX1 and 2 together play a more important role in HPAFII resistance to CAR T cells than COX2 by itself. This data is in line with other immunotherapy strategies such as checkpoint blockades combined with COX2 inhibitors [40]. Further studies are needed in order to explain the molecular events governing the success or failure of these combination therapies.

Since tMUC1-CAR T cell is new, we wished to confirm other published studies that combined CAR T cell treatment with standard of care chemotherapy drugs. Chemo and radiotherapy have primarily been used to make tumors leaky or to deplete lymphocytes to make new niches for CAR T cells prior to adoptive cell transfer. Results showed improved tumor regression and enhanced survival in metastatic melanoma [65, 66]. Moreover, chemotherapeutic drugs may be used to remove immunosuppressive T regulatory cells [67]. Data from this study clearly corroborate previous findings. Pre-treating resistant HPAFII cells with suboptimal dose of GEM, PTX or 5FU significantly enhanced CAR T cell cytotoxicity. To our surprise, PTX and GEM did not have the same effect on the CFPAC. In CFPAC, only 5FU was effective in enhancing CAR T cell treatment. Thus, we strongly believe that every tumor is different and responds differently to drugs and immunotherapy. These results provide a promising strategy for the potentiation of CAR T therapy in treating refractory PDA tumors.

Based on the *in vivo* orthotopic model of MiaPaCa2 tumors, tMUC1-CAR T cells could clearly reduce tumor burden but could not eradicate the tumor. To improve the treatment efficacy, several strategies can be utilized in the future, such as multiple injections of CAR T cells, combination with the other drugs mentioned above, and in the future using an immunocompetent model of PDA. Testing CAR T cells function in human MUC1 transgenic mouse model of spontaneous PDA, is currently under way by our group. In summary, our results provide promise for the use of tMUC1-CAR T cells against treatment refractory PDA in combination with inhibitors against IDO1, COX1/2 and Gal-9 or in combination with low-dose standard chemotherapy drugs.

## Acknowledgments

This study was supported by NIH 2 R01 CA135650-05A1 and NIH 1 R15 CA173668-01 grants. We thank Dr. Timothy Erick for editing the manuscript. We also thank our vivarium staff members for their assistance with training and animal care.

---

**Figure S1.**
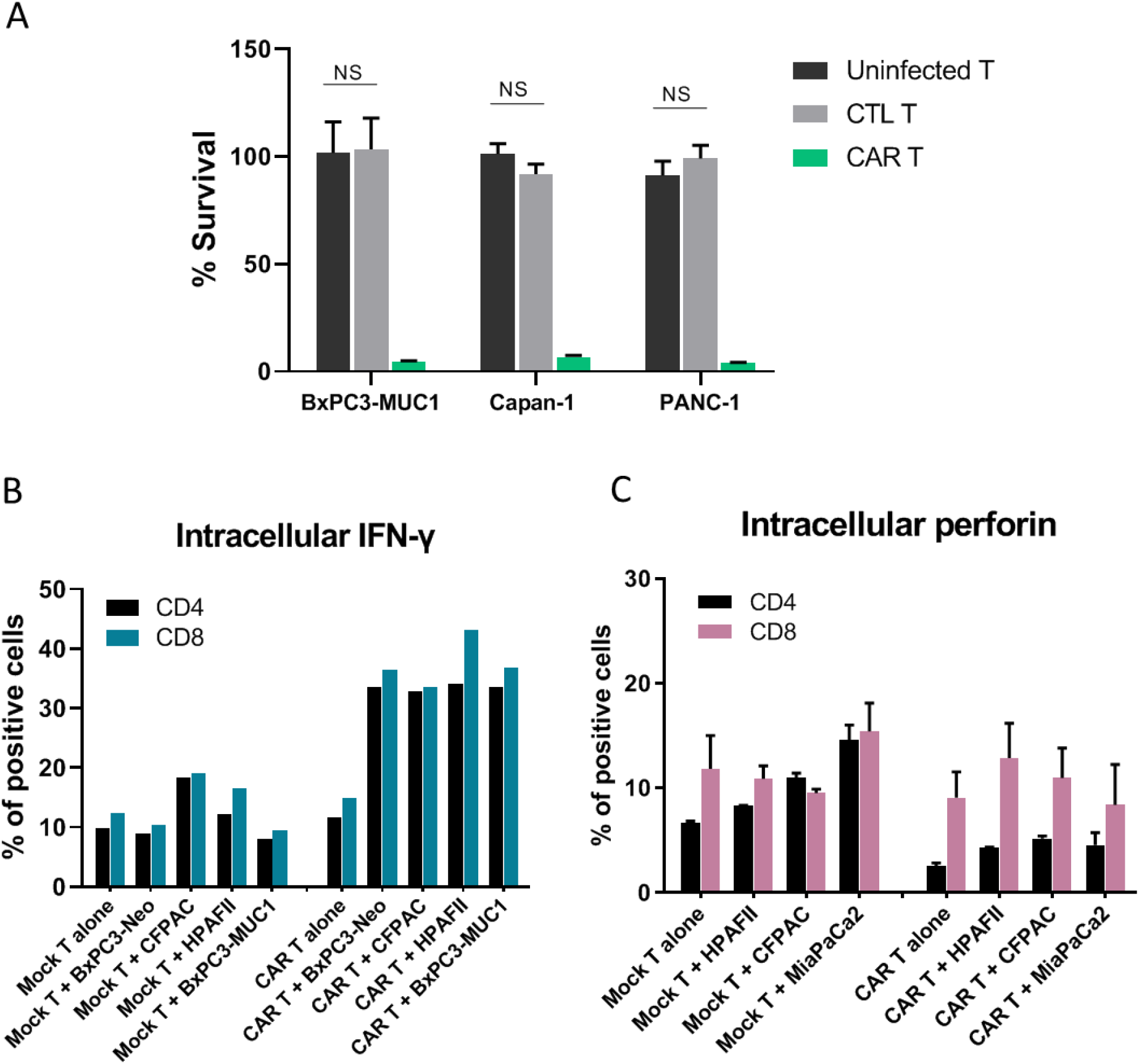
**A.** Similar activity of uninfected T and CTL T cells. Uninfected, CTL or CAR T cells were exposed to indicated PDA cell lines for 72 hrs at T:E 1:10, and survival level of PDA cells were measured using MTT assay. Data was normalized to media alone. Uninfected and CTL T show similar killing ability against PDA cells. Student’s t-test, NS P> 0.05. **B, C.** Intracellular level of IFN-γ **(B)** and perforin **(C)** in mock and CAR T cells before and after exposure to PDA cells for 24 hrs at T:E 1:10. Percentage of CD4^+^ and CD8^+^ CAR T cells that are positive for IFN-γ and perforin was measured by flowcytometry. Data suggests CAR T cells exposed to HPAFII and CFPAC resistant cells are not functionally impaired regarding the production of IFN-γ and perforin internally. Each graph is representative of three independent experiments.

**Figure S2.**
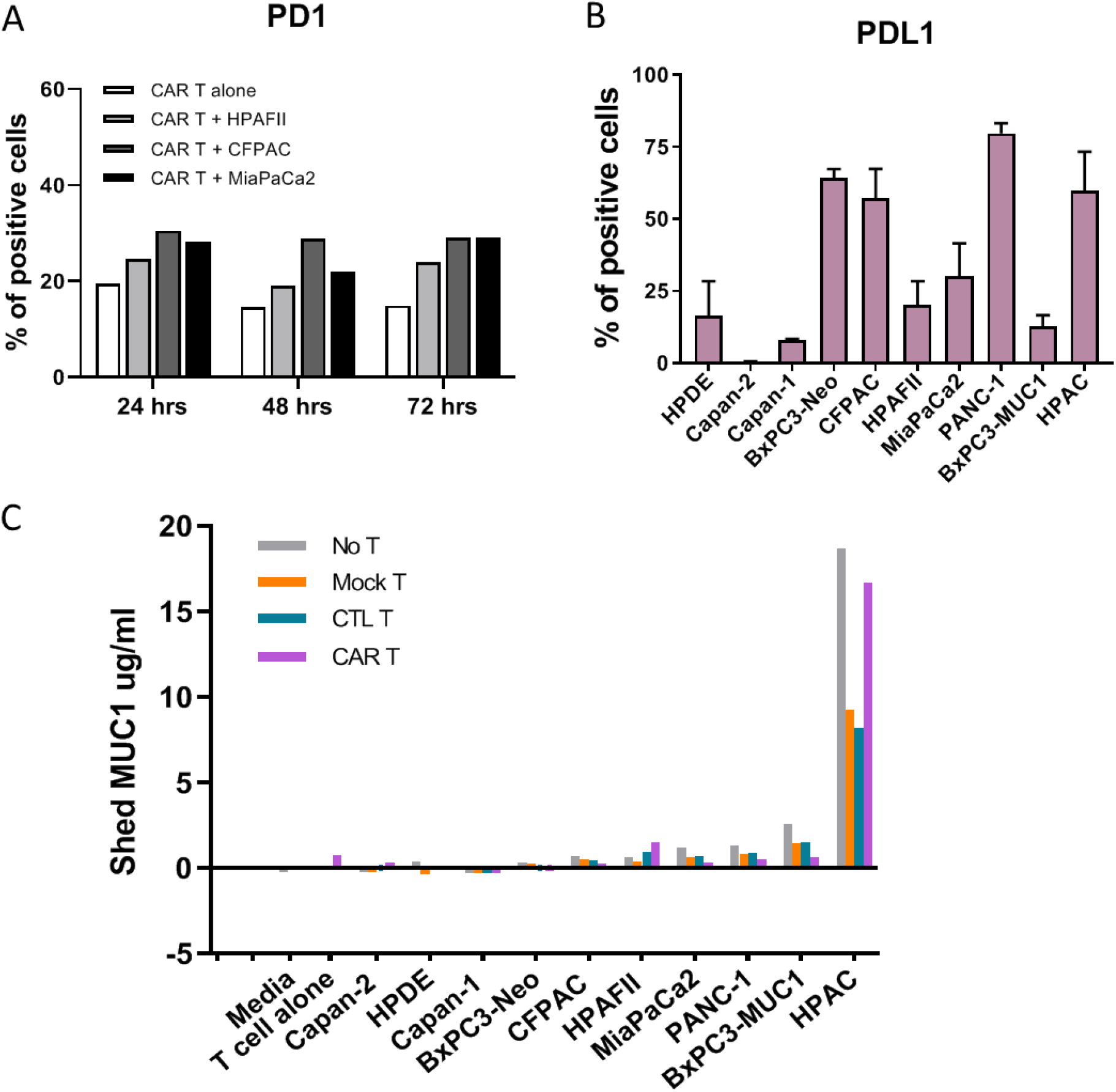
**A.** PD1 expression on CAR T cells before and after exposure to HPAFII, CFPAC and MiaPaCa2 cells for 24, 48 and 72hrs. Percentage of positive cells was measured using flowcytometry. CAR T cells exposed to PDA cells express higher level of PD1 compared to unexposed CAR T cells, however there was no significant difference in PD1 expression of CAR T cells when co-cultured with resistant (HPAFII and CFPAC) vs. sensitive cells (MiaPaca2). **B.** PDL1 expression in a panel of PDA cells measured using flowcytometry. There was no correlation between the PDL1 and resistance level in PDA cells. **C**. Level of shed MUC1 in the co-culture supernatant of PDA cells and T cells measured by ELISA. Most PDA cells and T cell alone do not shed noticeable amount of MUC1, however HPAC cells exceptionally released high amount of MUC1 into the media before and after exposure to CAR T cells (72 hrs). Data suggests shed MUC1 does not contribute to the immune resistance of HPAFII and CFPAC.

**Figure S3.**
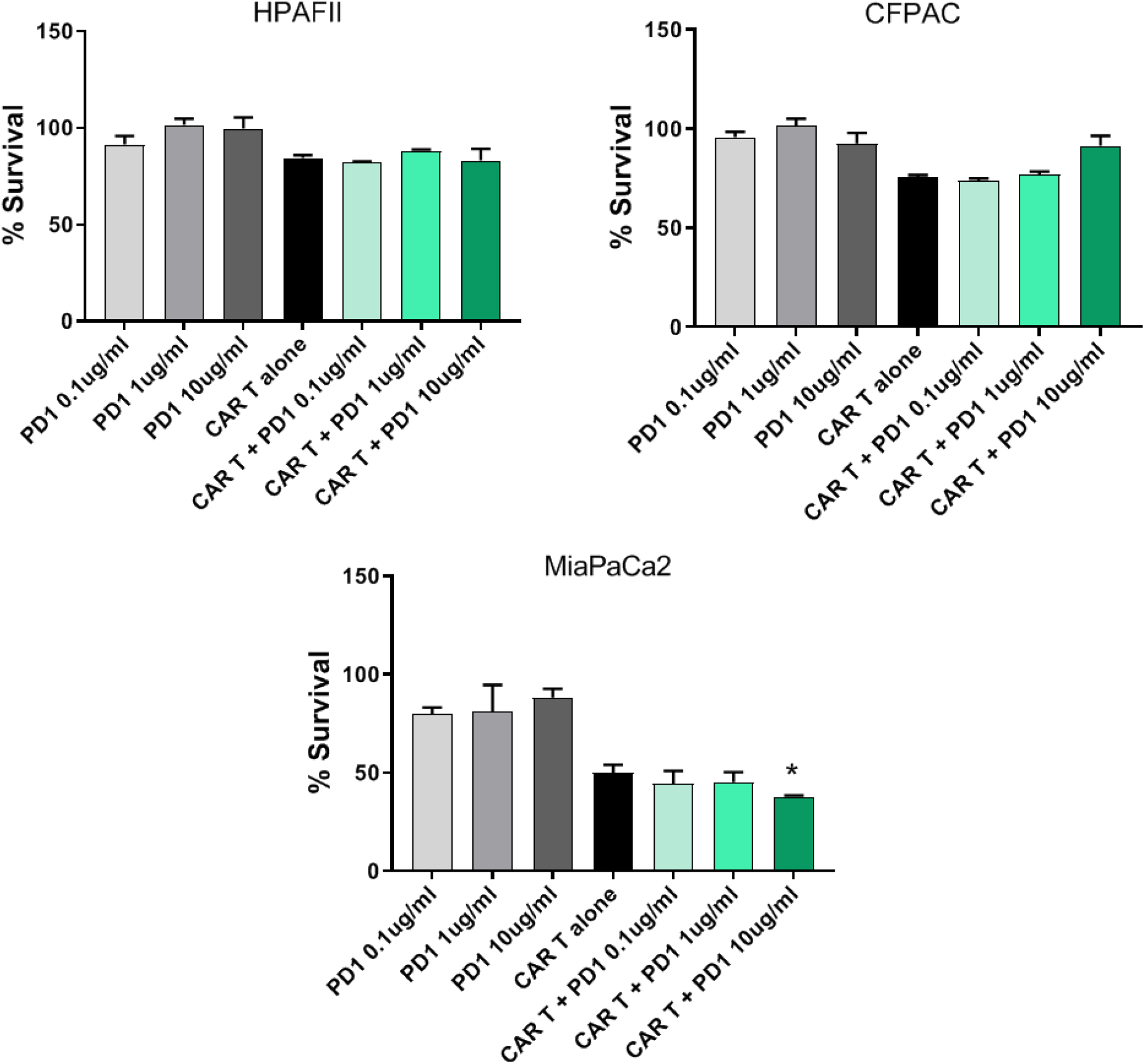
CAR T cell therapy in combination with anti-PD1 blocking antibody. Percentage survival of HPAFII, CFPAC and MiaPaCa2 PDA cells treated with PD1 blocking Ab alone, T cells alone, and combination, at 3 different concentrations of Ab. PDA cells were co-cultured with mock or CAR T cells, −/+ anti-PD1 blocking Ab for 72 hrs at T:E 1:10, and their survival level was measured using MTT assay. Percentage survival was normalized to mock T cells. Resistant cells killing by CAR T cells was not improved by adding anti-PD1 blocking Ab; while MiaPaCa2 cells killing was improved by the combination therapy. Unpaired Student’s t-test, comparing CAR T + PD1 group to CAR T alone group, * P=0.0404.

## References

1. Hidalgo, M., et al., Addressing the challenges of pancreatic cancer: Future directions for improving outcomes. Pancreatology, 2015. 15(1): p. 8–18.

2. Kamisawa, T., et al., Pancreatic cancer. Lancet, 2016. 388(10039): p. 73–85.

3. Yabar, C.S. and J.M. Winter, Pancreatic Cancer: A Review. Gastroenterol Clin North Am, 2016. 45(3): p. 429–45.

4. Swayden, M., J. Iovanna, and P. Soubeyran, Pancreatic cancer chemo-resistance is driven by tumor phenotype rather than tumor genotype. Heliyon, 2018. 4(12): p. e01055–e01055.

5. Srivastava, S. and S.R. Riddell, Engineering CAR-T cells: Design concepts. Trends in Immunology, 2015. 36(8): p. 494–502.

6. Kalos, M., et al., T Cells with Chimeric Antigen Receptors Have Potent Antitumor Effects and Can Establish Memory in Patients with Advanced Leukemia. Science Translational Medicine, 2011. 3(95): p. 95ra73.

7. Kochenderfer, J.N., et al., B-cell depletion and remissions of malignancy along with cytokine-associated toxicity in a clinical trial of anti-CD19 chimeric-antigen-receptor-transduced T cells. Blood, 2012. 119.

8. Rosenberg, S.A., et al., Treatment of Patients With Metastatic Melanoma With Autologous Tumor-Infiltrating Lymphocytes and Interleukin 2. Journal of the National Cancer Institute, 1994. 86(15): p. 1159–1166.

9. Yazdanifar, M., R. Zhou, and P. Mukherjee, Emerging immunotherapeutics in adenocarcinomas: A focus on CAR-T cells. Current Trends in Immunology, 2016. 17: p. 95–115.

10. Nath, S. and P. Mukherjee, MUC1: a multifaceted oncoprotein with a key role in cancer progression. Trends Mol Med, 2014. 20(6): p. 332–42.

11. Hollingsworth, M.A. and B.J. Swanson, Mucins in cancer: protection and control of the cell surface. Nat Rev Cancer, 2004. 4(1): p. 45–60.

12. Cheever, M.A., et al., The prioritization of cancer antigens: a national cancer institute pilot project for the acceleration of translational research. Clin Cancer Res, 2009. 15(17): p. 5323–37.

13. Levi, E., et al., MUC1 and MUC2 in pancreatic neoplasia. Journal of Clinical Pathology, 2004. 57(5): p. 456–462.

14. Chhieng, D.C., et al., MUC1 and MUC2 expression in pancreatic ductal carcinoma obtained by fine-needle aspiration. Cancer, 2003. 99(6): p. 365–71.

15. Schroeder, J.A., et al., MUC1 overexpression results in mammary gland tumorigenesis and prolonged alveolar differentiation. Oncogene, 2004. 23(34): p. 5739–47.

16. Sahraei, M., et al., MUC1 Regulates PDGFA Expression During Pancreatic Cancer Progression. Oncogene, 2012. 31(47): p. 4935–4945.

17. Beatty, G.L., et al., Mesothelin-specific chimeric antigen receptor mRNA-engineered T cells induce anti-tumor activity in solid malignancies. Cancer Immunol Res, 2014. 2(2): p. 112–20.

18. Hudecek, M., et al., Receptor affinity and extracellular domain modifications affect tumor recognition by ROR1-specific chimeric antigen receptor T cells. Clin Cancer Res, 2013. 19(12): p. 3153–64.

19. Berger, C., et al., Safety of targeting ROR1 in primates with chimeric antigen receptor-modified T cells. Cancer Immunol Res, 2015. 3(2): p. 206–16.

20. Maher, J., et al., Targeting of Tumor-Associated Glycoforms of MUC1 with CAR T Cells. Immunity, 2016. 45(5): p. 945–946.

21. Wilkie, S., et al., Retargeting of human T cells to tumor-associated MUC1: the evolution of a chimeric antigen receptor. J Immunol, 2008. 180(7): p. 4901–9.

22. Wilkie, S., et al., Dual targeting of ErbB2 and MUC1 in breast cancer using chimeric antigen receptors engineered to provide complementary signaling. J Clin Immunol, 2012. 32(5): p. 1059–70.

23. Anurathapan, U., et al., Kinetics of Tumor Destruction by Chimeric Antigen Receptor-modified T Cells. Mol Ther, 2014. 22(3): p. 623–633.

24. Posey, A.D., Jr., et al., Engineered CAR T Cells Targeting the Cancer-Associated Tn-Glycoform of the Membrane Mucin MUC1 Control Adenocarcinoma. Immunity, 2016. 44(6): p. 1444–54.

25. You, F., et al., Phase 1 clinical trial demonstrated that MUC1 positive metastatic seminal vesicle cancer can be effectively eradicated by modified Anti-MUC1 chimeric antigen receptor transduced T cells. Science China Life Sciences, 2016. 59(4): p. 386–397.

26. Roy, L.D., et al., A tumor specific antibody to aid breast cancer screening in women with dense breast tissue. Genes & Cancer, 2017. 8(3–4): p. 536–549.

27. Moore, L.J., et al., Antibody-Guided In Vivo Imaging for Early Detection of Mammary Gland Tumors. Transl Oncol, 2016. 9(4): p. 295–305.

28. Zhou, R., et al., A novel association of neuropilin-1 and MUC1 in pancreatic ductal adenocarcinoma: role in induction of VEGF signaling and angiogenesis. Oncogene, 2016. 35(43): p. 5608–5618.

29. Curry, J.M., et al., The use of a novel MUC1 antibody to identify cancer stem cells and circulating MUC1 in mice and patients with pancreatic cancer. J Surg Oncol, 2013. 107(7): p. 713–22.

30. Roy, L.D., et al., Early detection of breast cancer using a unique tumor specific antibody. J Clin Oncol, 2015. 33: p. abstr e22153.

31. Wu, S.-T., et al., Treatment of pancreatic ductal adenocarcinoma with tumor antigen specific-targeted delivery of paclitaxel loaded PLGA nanoparticles. BMC Cancer, 2018. 18(1): p. 457.

32. Dréau, D., et al., Mucin-1-Antibody-Conjugated Mesoporous Silica Nanoparticles for Selective Breast Cancer Detection in a Mucin-1 Transgenic Murine Mouse Model. Journal of biomedical nanotechnology, 2016. 12(12): p. 2172–2184.

33. Silvestri, I., et al., A Perspective of Immunotherapy for Prostate Cancer. Cancers (Basel), 2016. 8(7): p. E64

34. Morgan, R.A., et al., Case Report of a Serious Adverse Event Following the Administration of T Cells Transduced With a Chimeric Antigen Receptor Recognizing ERBB2. Molecular Therapy, 2010. 18(4): p. 843–851.

35. Delitto, D., S.M. Wallet, and S.J. Hughes, Targeting tumor tolerance: A new hope for pancreatic cancer therapy? Pharmacol Ther, 2016. 166: p. 9–29.

36. Whilding, L.M., et al., Targeting of Aberrant alphavbeta6 Integrin Expression in Solid Tumors Using Chimeric Antigen Receptor-Engineered T Cells. Mol Ther, 2017. 25(1): p. 259–273.

37. Waterhouse, N.J., et al., Cytotoxic T lymphocyte-induced killing in the absence of granzymes A and B is unique and distinct from both apoptosis and perforin-dependent lysis. J Cell Biol, 2006. 173(1): p. 133–44.

38. Zhou, R., et al., A novel association of neuropilin-1 and MUC1 in pancreatic ductal adenocarcinoma: role in induction of VEGF signaling and angiogenesis. Oncogene, 2016: p. Epub ahead of print.

39. Chan, A.K., et al., Soluble MUC1 secreted by human epithelial cancer cells mediates immune suppression by blocking T-cell activation. Int J Cancer, 1999. 82(5): p. 721–6.

40. Zelenay, S., et al., Cyclooxygenase-Dependent Tumor Growth through Evasion of Immunity. Cell, 2015. 162(6): p. 1257–70.

41. Mukherjee, P., et al., Progression of Pancreatic Adenocarcinoma Is Significantly Impeded with a Combination of Vaccine and COX-2 Inhibition(). Journal of immunology (Baltimore, Md.: 1950), 2009. 182(1): p. 216–224.

42. Fujihara, S., et al., Galectin-9 in cancer therapy. Recent Pat Endocr Metab Immune Drug Discov, 2013. 7(2): p. 130–7.

43. Zhou, R., et al., CAR T Cells Targeting the Tumor MUC1 Glycoprotein Reduce Triple-Negative Breast Cancer Growth. Front. Immunol., 2019.

44. John, L.B., et al., Anti-PD-1 antibody therapy potently enhances the eradication of established tumors by gene-modified T cells. Clin Cancer Res, 2013. 19(20): p. 5636–46.

45. Zhou, R., et al., The use of tMUC1 highly specific chimeric antigen receptor-redirected T cells for the eradication of triple negative breast cancer. The Journal of Immunology, 2017. 198(1 Supplement): p. 198.10.

46. Koyama, S., et al., Adaptive resistance to therapeutic PD-1 blockade is associated with upregulation of alternative immune checkpoints. Nature communications, 2016. 7: p. 10501–10501.

47. van Baren, N. and B.J. Van den Eynde, Tryptophan-degrading enzymes in tumoral immune resistance. Front Immunol, 2015. 6: p. 34.

48. Pilotte, L., et al., Reversal of tumoral immune resistance by inhibition of tryptophan 2,3-dioxygenase. Proc Natl Acad Sci U S A, 2012. 109(7): p. 2497–502.

49. Frumento, G., et al., Tryptophan-derived catabolites are responsible for inhibition of T and natural killer cell proliferation induced by indoleamine 2,3-dioxygenase. The Journal of experimental medicine, 2002. 196(4): p. 459–468.

50. Hwu, P., et al., Indoleamine 2,3-dioxygenase production by human dendritic cells results in the inhibition of T cell proliferation. J Immunol, 2000. 164(7): p. 3596–9.

51. Terness, P., et al., Inhibition of allogeneic T cell proliferation by indoleamine 2,3-dioxygenase-expressing dendritic cells: mediation of suppression by tryptophan metabolites. J Exp Med, 2002. 196(4): p. 447–57.

52. Munn, D.H., et al., Inhibition of T cell proliferation by macrophage tryptophan catabolism. J Exp Med, 1999. 189(9): p. 1363–72.

53. Nagahara, K., et al., Galectin-9 Increases Tim-3^+^ Dendritic Cells and CD8^+^ T Cells and Enhances Antitumor Immunity via Galectin-9-Tim-3 Interactions. The Journal of Immunology, 2008. 181(11): p. 7660.

54. Golden-Mason, L., et al., Galectin-9 Functionally Impairs Natural Killer Cells in Humans and Mice. Journal of Virology, 2013. 87(9): p. 4835.

55. Goncalves Silva, I., et al., The Tim-3-galectin-9 Secretory Pathway is Involved in the Immune Escape of Human Acute Myeloid Leukemia Cells. EBioMedicine, 2017. 22: p. 44–57.

56. He, Y., et al., TIM-3, a promising target for cancer immunotherapy. OncoTargets and therapy, 2018. 11: p. 7005–7009.

57. Zhu, C., et al., The Tim-3 ligand galectin-9 negatively regulates T helper type 1 immunity. Nat Immunol, 2005. 6(12): p. 1245–52.

58. Kang, C.-W., et al., Apoptosis of tumor infiltrating effector TIM-3+CD8+ T cells in colon cancer. Scientific reports, 2015. 5: p. 15659–15659.

59. Gleason, M.K., et al., Tim-3 is an inducible human natural killer cell receptor that enhances interferon gamma production in response to galectin-9. Blood, 2012. 119(13): p. 3064–3072.

60. Galore-Haskel, G., et al., A novel immune resistance mechanism of melanoma cells controlled by the ADAR1 enzyme. Oncotarget, 2015. 6(30): p. 28999–9015.

61. Ishizuka, J.J., et al., Loss of ADAR1 in tumours overcomes resistance to immune checkpoint blockade. Nature, 2019. 565(7737): p. 43–48.

62. Pang, L.Y., E.A. Hurst, and D.J. Argyle, Cyclooxygenase-2: A Role in Cancer Stem Cell Survival and Repopulation of Cancer Cells during Therapy. Stem cells international, 2016. 2016: p. 2048731–2048731.

63. Kalinski, P., Regulation of Immune Responses by Prostaglandin E_2_. The Journal of Immunology, 2012. 188(1): p. 21.

64. Sobolewski, C., et al., The Role of Cyclooxygenase-2 in Cell Proliferation and Cell Death in Human Malignancies. International Journal of Cell Biology, 2010. 2010.

65. Koike, N., S. Pilon-Thomas, and J.J. Mule, Nonmyeloablative chemotherapy followed by T-cell adoptive transfer and dendritic cell-based vaccination results in rejection of established melanoma. J Immunother, 2008. 31(4): p. 402–12.

66. Dudley, M.E., et al., Adoptive cell transfer therapy following non-myeloablative but lymphodepleting chemotherapy for the treatment of patients with refractory metastatic melanoma. J Clin Oncol, 2005. 23(10): p. 2346–57.

67. Wang, L., et al., Chimeric antigen receptor T cell therapy and other therapeutics for malignancies: Combination and opportunity. Int Immunopharmacol, 2019. 70: p. 498–503.

